# Filaggrin deficient mice have a lower threshold for cutaneous allergen sensitization but do not develop spontaneous skin inflammation or atopy

**DOI:** 10.1101/2020.09.11.293688

**Authors:** Lina Muhandes, Martin Pippel, Maria Chapsa, Rayk Behrendt, Yan Ge, Andreas Dahl, Buqing Yi, Alexander Dalpke, Sylke Winkler, Michael Hiller, Stefan Beissert, Rolf Jessberger, Padraic G. Fallon, Axel Roers

## Abstract

Defects of filaggrin (FLG) compromise epidermal barrier function and represent an important known genetic risk factor for atopic dermatitis (AD), but also for systemic atopy, including allergic sensitization and asthma. The flaky tail mouse model, widely used to address mechanisms of atopy induction by barrier-defective skin, harbors two mutations that affect the skin barrier, the mutation *Flg*^*ft*^, resulting in near-complete loss of FLG expression, and the matted mutation inactivating transmembrane protein 79 (Tmem79). Upon separation of the two mutant loci, which are closely linked on chromosome 3, mice defective only for Tmem79 featured pronounced dermatitis and systemic atopy. Upon extensive backcross to BALB/c, also *Flg*^*ft/ft*^ mice (assumed to be wild type for *Tmem79*), developed AD-like dermatitis and reproduced the human ‘atopic march’, with high IgE levels and spontaneous asthma, suggesting a key role for functional Flg in protection from atopy also in mice. In contrast, BALB/c mice congenic for a targeted *Flg* knock out mutation did not develop skin inflammation or atopy. To resolve this discrepancy, we generated Flg-deficient mice on a pure BALB/c background by inactivating the *Flg* gene in BALB/c embryos. These animals feature an ichthyosis phenotype, but do not develop spontaneous dermatitis or systemic atopy. We sequenced the genome of the atopic *Flg*^*ft*^ BALB/c congenics and discovered that they were unexpectedly homozygous for the atopy-causing *Tmem79*^matted^ mutation. In summary, we show that Flg-deficiency does not cause atopy in mice. This finding is in line with lack of atopic disease in a fraction of Ichthyosis vulgaris patients carrying two FLG null alleles. However, absence of FLG may promote and modulate dermatitis caused by other genetic barrier defects, as skin inflammation in *Tmem79*^*ma*/ma^*Flg*^*ft/ft*^ BALB/c congenics is qualitatively different compared to *Tmem79*^*ma*/ma^ mice.

## Introduction

Atopic dermatitis (AD) is a multifactorial chronic inflammatory skin disorder representing a major socio-economic burden with a steady rise in incidence (reviewed in (Bieber, 2008; Weidinger and Novak, 2016)). AD currently affects up to one third of children in ‘Western’ countries. Childhood AD strongly predisposes to development of type I allergies and asthma later in life, a phenomenon addressed as the ‘atopic march’. All these atopic conditions share a complex genetic predisposition (reviewed in (Otsuka et al., 2017)). Traditionally, defects intrinsic to the immune system resulting in dysregulation of T cell polarization, including dominance of T_H_2 responses, were thought to be the primary cause of AD, type I allergies and asthma. Over the past two decades, however, the discovery of structural defects of the epidermal skin barrier as a cause of systemic immune dysregulation, leading not only to dermatitis, but also increased frequencies of allergic sensitization, high IgE and asthma, shifted this paradigm (Palmer et al., 2006; Walley et al., 2001). Barrier-defective skin allows abnormal penetration of environmental antigen. Presentation of these antigens occurs in the context of an abnormal immune milieu, as stressed skin cells, including keratinocytes, insufficiently protected against environmental hazards, activate innate immune responses. The stressed cells chronically release cytokines that drive the pathogenic Th2 bias and prominent immunoglobulin class switch to IgE that characterizes atopy (Dickel et al., 2010; Kakkar et al., 2012; Roan et al., 2019; Savinko et al., 2012).

Integrity of the epithelial skin barrier depends on continuous proliferative activity of the epidermis maintaining supply of anuclear corneocytes to renew the outermost layer of the skin, the Stratum corneum, which is the most critical component of the skin barrier (reviewed in (Matsui and Amagai, 2015)). The cornified layer is composed of corneocytes tightly connected by specialized desmosomes. The corneocyte is devoid of organelles and essentially consist of a matrix of compacted keratin filaments surrounded by a proteinaceous shell, the cornified envelope, that replaces the plasma membrane. The space between the corneocytes is filled by a complex mixture of lipids, making the cornified layer watertight. While terminally differentiating into corneocytes, keratinocytes express the S100-fused type protein (SFTP) filament aggregating protein (filaggrin, FLG). Full-length profilaggrin, one of the largest human proteins (400 kDa), is proteolytically cleaved into FLG monomers which bind to and bundle keratin intermediate filaments, thereby condensing the cytoskeleton and compacting the cell (Sandilands et al., 2009). FLG is further cleaved into hygroscopic amino acids which represent a ‘natural moisturizing factor’ (NMF) ensuring appropriate water content but also acidification of the cornified layer (Sandilands et al., 2009).

Genetic defects of FLG were identified as the cause of ichthyosis vulgaris (IV), a common skin disorder characterized by dry and scaly skin (Smith et al., 2006). The mode of inheritance is semi-dominant with heterozygous loss-of-function mutations causing a mild phenotype (Smith et al., 2006). Most patients with severe IV also develop atopy and suffer from AD, allergic sensitizations and asthma (Palmer et al., 2006). Conversely, sequencing of the FLG gene in cohorts of patients with asthma and AD revealed 1 or 2 null alleles in 23% of cases, establishing genetic FLG deficiency as a major factor predisposing for AD and asthma associated with AD (Palmer et al., 2006; Sandilands et al., 2007)(reviewed in (Irvine et al., 2011)). Strong association of FLG mutations were also reported for pollen and food allergies (reviewed in (Irvine et al., 2011)).

Given this key role of barrier-defects in atopic disease, animal models of FLG deficiency are of great interest. The flaky tail mouse (Presland et al., 2000) carries genetic defects of two genes involved in epidermal barrier function, the *Flg*^*ft*^ mutation, a 1 bp deletion (*Flg* 5303delA), resulting in hypomorphic FLG expression (Fallon et al., 2009), and a deleterious mutation in the gene encoding transmembrane protein 79 (Saunders et al., 2013). The *Tmem79*^*ma*^ mutation (p.Y280*) results in impaired cornification with matted fur. The two genes are closely linked on mouse chromosome 3 (Saunders et al., 2013). With this dual epidermal barrier impairment, flaky tail mice develop dry and scaly skin, eczema and systemic atopy with increased serum IgE (Fallon et al., 2009; Lane, 1972; Presland et al., 2000).

Addressing the individual contributions of the two genetic defects to the atopic phenotype, Saunders et al. separated the *Tmem79*^*ma*^ mutation from the *Flg*^*ft*^ allele and showed that isolated loss of Tmem79 results in eczema and increased serum IgE(Saunders et al., 2013). Targeted complete inactivation of *Flg* in C57BL/6 resulted in the IV-like phenotype as expected, however, this mutation did not trigger atopy even after backcrossing to BALB/c (Kawasaki et al., 2012). In striking contrast to this finding, Saunders et al. reported that extensive backcrossing of the *Flg*^*ft*^ mutation to BALB/c yielded mice which developed severe skin inflammation with important features of AD, including elevated serum IgE levels, eosinophilia and accumulation of IL-5-producing ILC2s. (Saunders et al., 2016). Most importantly, these *Flg*^*ft*^ BALB/c congenics uniformly show high levels of total IgE and develop spontaneous asthma, thus reproducing the ‘atopic march’ observed in many human patients (Saunders et al., 2016). These findings established *Flg*^ft/ft^ BALB/c congenic mice as a highly relevant model for human atopy associated with skin barrier defects.

The human *FLG* and mouse *Flg* genes are encoded within a large cluster of genes functioning in formation and maintenance of the skin barrier, the epidermal differentiation cluster (EDC) and are in strong linkage disequilibrium with the entire EDC. To determine whether genetic variability within the EDC and adjacent regions may account for the phenotypic difference between the *Flg*^ft/ft^ BALB/c congenics (Saunders et al., 2016) and the Flg knock out mice (Kawasaki et al., 2012), we generated Flg-deficient mice on a pure BALB/c background and analysed them for skin phenotype and systemic atopy. We also performed whole genome sequencing for comparison of the *Flg*^ft/ft^ BALB/c congenics and our new BALB/c *Flg*^*-/-*^ line.

## Results

### Generation of BALB/c *Flg*^*-/-*^ mice

For inactivation of the *Flg* gene, two guide RNA sequences were chosen. Guide 1 targeted the upstream, non-repetitive portion of exon 3, whereas guide 2 bound each of the repeat regions of exon 3 (Fig. 1A). The guides were microinjected into BALB/c zygotes and mutant offspring were identified by PCR (Fig S1). We found three mutant *Flg* alleles that harbored large deletions of the upstream non-repetitive region plus at least 1 *Flg* repeat as determined by PCR (Fig S1 B and C). The mutations were bred to homozygosity and total tail and ear skin samples from 8 week-old mice were analyzed by Western blot, demonstrating complete absence of both, full-length profilaggrin and monomeric filaggrin protein (Fig 1B). This result was confirmed by immunofluorescence analysis with no filaggrin protein expressed in skin of *Flg*^*-/-*^mice (Fig 1C). Mice homozygous for allele 1 (Fig. S1C) were used in subsequent experiments. Long read single molecule real-time (SMRT) sequencing (PacBio) revealed that this line carried a large (8122 bp) out-of-frame deletion (Fig. 1A). In sum, we successfully generated FLG-deficient mice on a pure BALB/c background.

**Figure 1.**
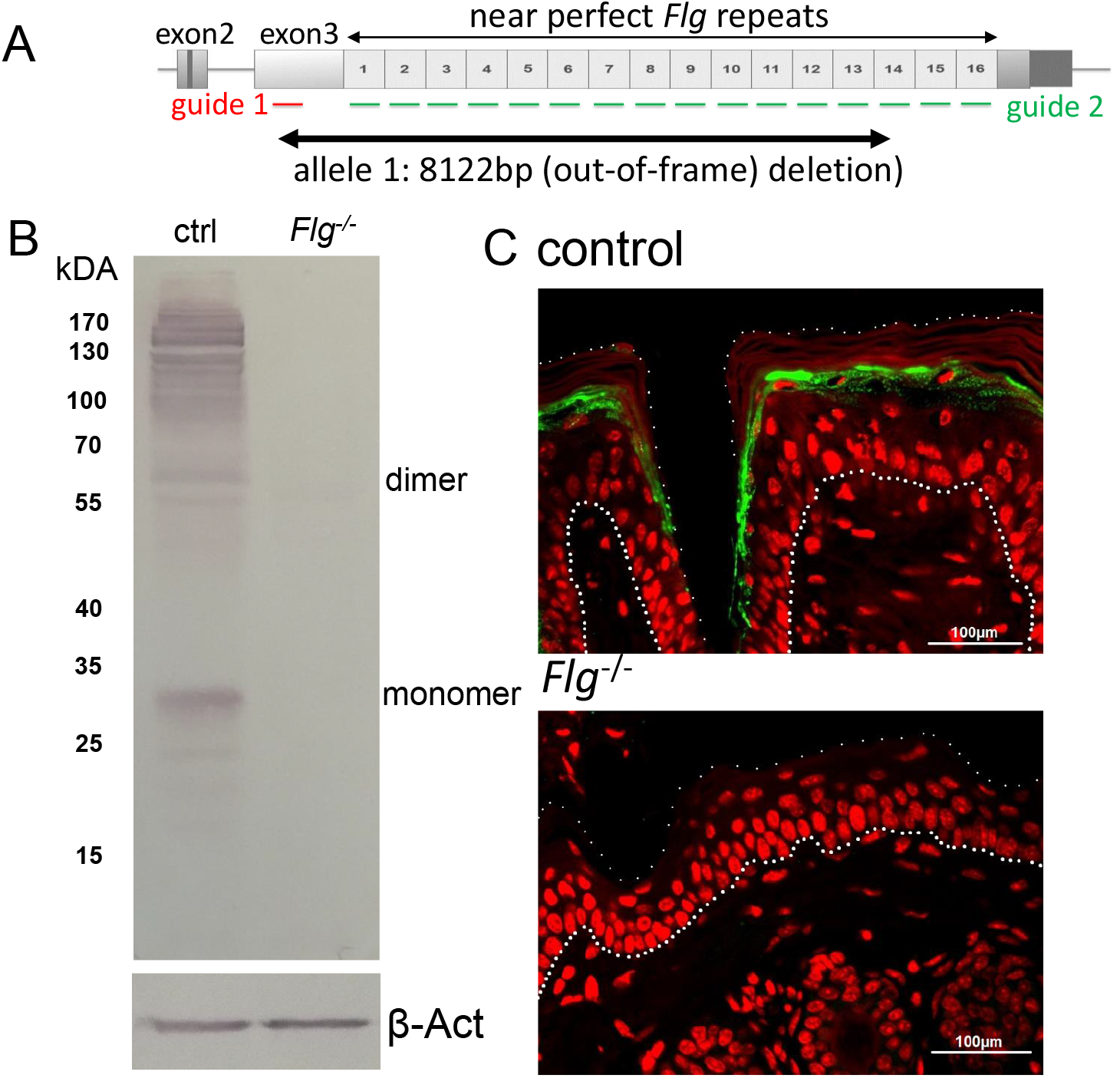
Generation of Flg-/-mice on a pure BALB/c background. A) Strategy for CRISPR/Cas9-mediated targeting of the *Flg* gene. While guide 1 binds a single site in the upstream, non-repetitive region of exon 3, guide 2 binds to each of the near perfect *Flg* repeats. Among others we obtained mutant ‘allele1’ that was bred to homozygosity to obtain *Flg*^*-/-*^ mice used in subsequent experiments. Long read single molecule real-time sequencing revealed that allele1 contained a 8122bp (out-of-frame) deletion. B) Expression of FLG in ear skin as determined by western blot analysis of ear skin from 8 week-old mice. Expected size for full-length profilaggrin is approximately 500 kDa and approximately 30 kDa for the FLG monomer. Representative of 8 animals. C) Immunofluorescence analysis FLG expression (green) in tail skin of 8-week-old control (*Flg*^wt/-^) and *Flg*^*-/-*^mice. Nuclei counterstained with DAPI (red). Representative images from 4 mice of each genotype. Dotted lines mark basement membrane and the border of epidermis to cornified layer. Scale bar 100 µM.

### BALB/c *Flg*^*-/-*^ mice develop an ichthyosis vulgaris-like phenotype

While normal immediately after birth, homozygous *Flg*-deficient pups became conspicuous few days later for their dry and scaly skin, hyperlinearity and annular constrictions of the tail skin (Fig. 2A and B). This phenotype ameliorated over the first few weeks post-partum and the skin appeared macroscopically normal at latest by the age of 4 weeks (not shown). There was no evidence of pruritus and the fur of adult mice was indistinguishable from that of control littermates. Hematoxylin and eosin staining of skin sections revealed marked reduction of keratohyalin granules in the upper *Flg*^*-/-*^ epidermis (Stratum granulosum), consistent with lack of Flg as the major constituent of these granules and reproducing a histological hallmark of IV (Sybert et al., 1985). Apart from loss of keratohyalin granules, *Flg*^*-/-*^ epidermis appeared normal, as did the dermis with no evidence of inflammatory infiltration. While *Flg*^*-/-*^ ears were invariably smaller (not shown) and thicker (Fig. 2C) as had been described for the flaky tail strain (Lane, 1972), no overt histological changes were observed in ear skin. Transepidermal water loss (TEWL) was unchanged compared to controls in younger *Flg*^*-/-*^ mice (8-10 weeks), but was clearly increased in older filaggrin deficient (16-18 weeks) animals compared to age-matched controls (Fig. 2D).

**Figure 2.**
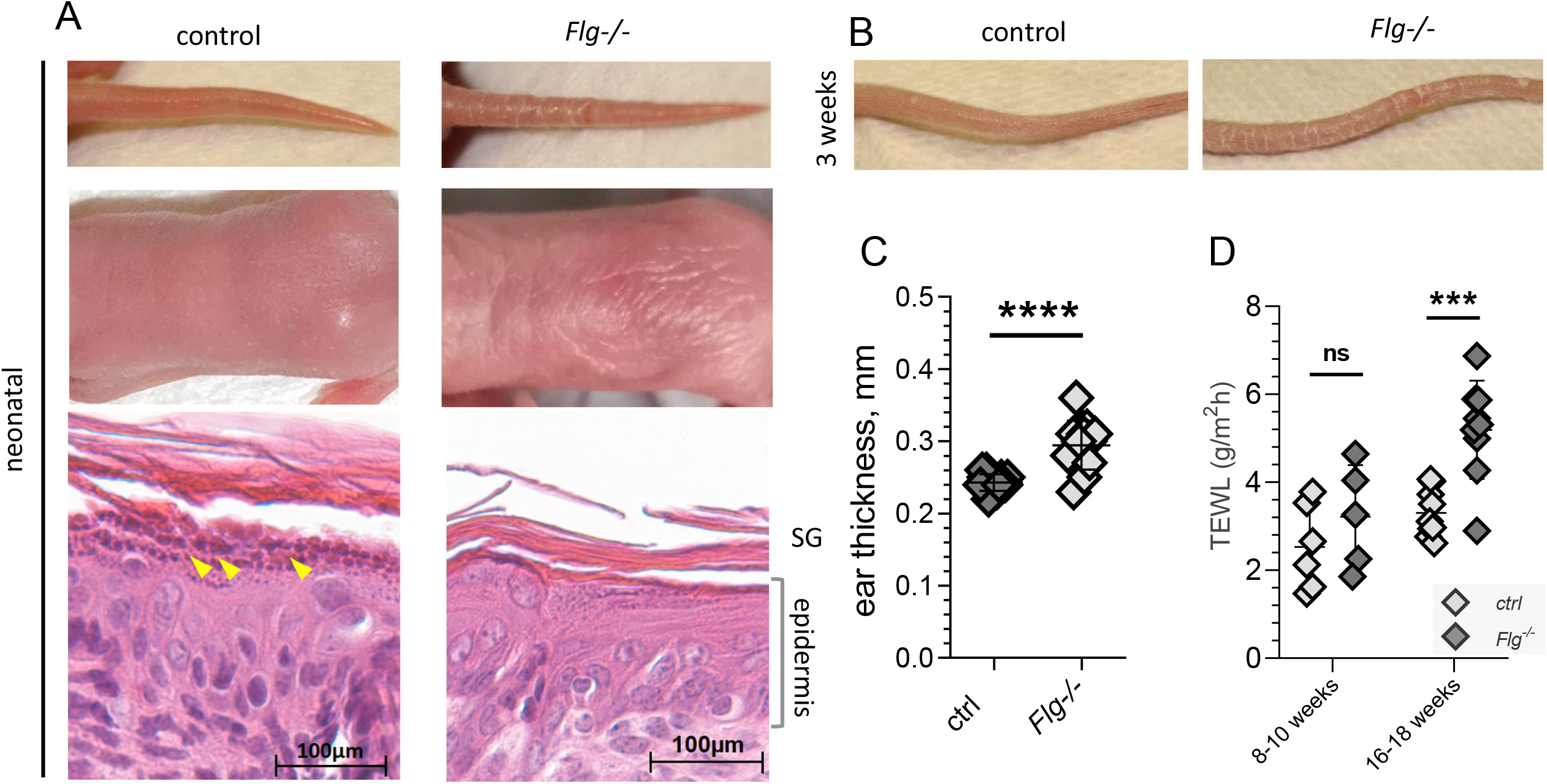
Skin phenotype of BALB/c *Flg*^*-/-*^ mice. A) Representative macroscopic and microscopic images of neonatal (d4) BALB/c control and *Flg*^*-/-*^ ear skin. Lower panels: H&E staining of epidermis. Insets from larger images shown in Fig. S2. Arrowheads denote keratohyalin granules of control skin. Images representative of at least 10 mice. Scale bar 100 µM. B) Representative macroscopic images of 3 week-old BALB/c control and *Flg*^*-/-*^ tails. C) Ear thickness of 8 week-old BALB/c littermate control (n=14) and *Flg*^*-/-*^ (n=13), male and female both groups. Means ± SD, ****p< 0.0001. D) Transepidermal water loss (TEWL) of shaved BALB/c control and *Flg*^*-/-*^ back skin. Means ± SD, ***p< 0.001, ns, not significant.

Collectively, we find that loss of FLG on a pure BALB/c background results in dry and scaly skin in neonates. In adult mice, skin and fur are macroscopically normal. Increased TEWL occurs in older BALB/c *Flg*^*-/-*^ animals, but not in younger adults, indicative of a defect in the skin barrier in adult *Flg*^*-/-*^ mice.

### BALB/c *Flg*^*-/-*^ skin features mild immune activation and altered microbiome

Skin barrier defects can result in systemic immune dysregulation through cytokine release of stressed epidermal cells that are insufficiently protected against environmental hazards. In order to test, whether the loss of physiological Flg expression in the suprabasal epidermis leads to detectable stress responses of cells of lower epidermal strata, we performed RNA sequencing of FACS-sorted CD45-negative ITGA6 (CD49f)^+^ basal keratinocytes of young adult control and BALB/c *Flg*^*-/-*^ mice (Fig. S3A). Irrespective of genotype, transcripts known to be expressed by basal layer keratinocytes were abundant (keratin 1, 5 and 10) while genes expressed only in suprabasal cells, including *Flg, Ivl* (involucrin), *lor* (loricrin), *Tgm* (transglutaminase) *1, 3* and *5* were not detected, validating the experiment. In accordance with the moderate changes observed macroscopically and histologically, only few genes showed a significant up-regulation and we observed this only in female mice, while males did not feature significantly altered gene expression. Among the genes upregulated in females were several genes encoding components of the MHC class II antigen presentation machinery, including *H2-Aa, H2-Ab1, H2-Ea, CD74* (encoding invariant chain) and *H2-DMb1* (Fig. 3A). qRT PCR analysis for *Cd74* and *H2-Aa* were found upregulated (although not statistically significant for individual genes) (Fig. 3B). Flow cytometric analysis of skin cell suspensions demonstrated a doubling of MHC class II expressing keratinocytes in female mice (Fig. 3C). Patches of brightly MHC II-positive keratinocytes were recently identified as critical structures in antimicrobial T cell responses in the skin (Tamoutounour et al., 2019). We also found a robust upregulation of the Retnla gene encoding restin-like molecule alpha (RELMα) in sorted female, but not male *Flg*^*-/-*^ basal keratinocytes (Figure 3A and B). RELMα is a marker of M2 macrophage differentiation and was also demonstrated to function as an important keratinocyte antibacterial effector protein (Harris et al., 2019). While expression of IL-33 was not different between BALB/c *Flg*^*-/-*^ mice and controls (not shown), IL-1β mRNA levels, while unchanged in young adults, were clearly increased in 20 weeks-old mice (Figure 3D).

**Figure 3.**
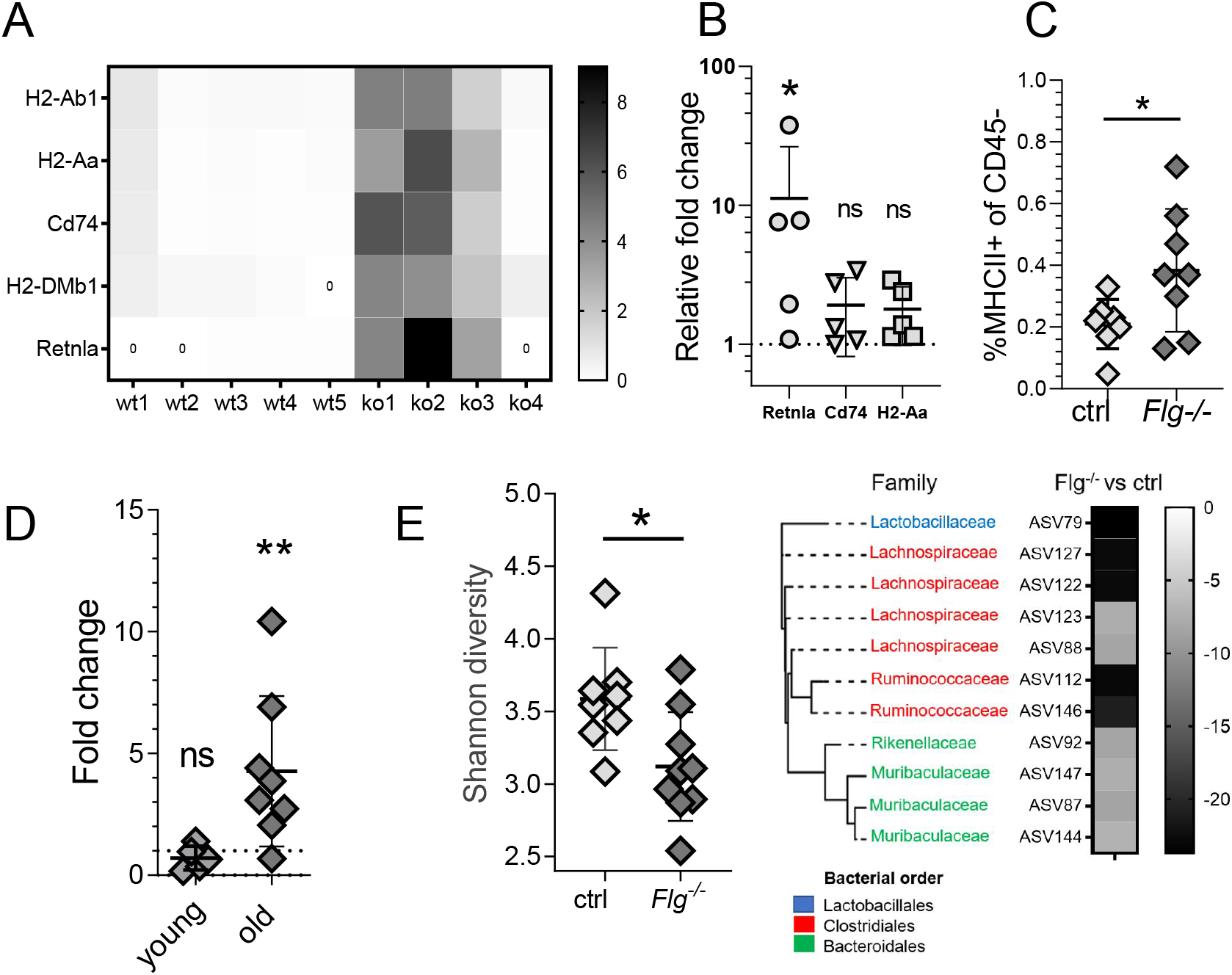
Alteration of keratinocyte gene expression and composition of microbiota of *Flg*^*-/-*^ skin. A) RNA sequencing analysis of basal CD49f^+^ keratinocytes (negative for CD45, see also Fig S3), sorted from skin cell suspensions of four 8-12 week-old female *Flg*^*-/-*^ mice and 5 littermate controls. Heat map represents expression of genes involved in antigen presentation and the *Retnla* gene (relative expression, individual read counts corrected for the mean expression of the respective gene), p<0.05 for all genes. B) Validation of results in A) by qRT-PCR for selected genes on RNA extracted from basal keratinocytes sorted as in A). Fold change for individual female *Flg*^*-/-*^ mice displayed as fold change compared to mean of 6 controls (set to 1). Means ± SD, *p<0.05, ns, not significant. C) Frequency of CD45-negative epidermal cells expressing MHCII as determined by flow cytometric analysis of skin cell suspensions from 8-10 week-old *Flg*^*-/-*^ and control females. n=8 both groups. Means ± SD, p< 0.05. D) Relative fold change of IL-1β mRNA in total ear skin RNA from young adult (8 wk-old)) and older (20 wk-old) *Flg*^*-/-*^ BALB/c mice compared to controls. Means ± SD, n=8 both groups,**p<0.01, ns, not significant. E) Composition of the skin microbiome of 8 week-old female BALB/c *Flg*^*-/-*^ and control co-housed females. Left: comparison of Shannon diversity index. Right: Phylogeny and abundance of taxa that were significantly depleted in *Flg*^*-/-*^ mice. Branch label colors are indicative of the bacterial order that the respective amplicon sequence variant (ASV) was assigned to. Heatmap displays log2 fold change of each ASV for the comparison of *Flg*^*-/-*^ versus control.

To determine whether absence of Flg affects the skin microbiome, we swabbed cheek skin of 8 week-old female BALB/c *Flg*^*-/-*^ and control mice co-housed in the same cages and performed 16s ribosomal RNA sequencing. As expected for mouse skin, *Bacteroides, Proteobacteria* and *Firmicutes* were the dominant phyla without major changes of their abundance in mutants versus controls (Fig. S3B). BALB/c *Flg*^*-/-*^ mice showed a reduction of microbial diversity compared to control as indicated by Shannon diversity index (Fig. 3E). Several taxa of the families *Muribaculaceae*,previously known as S24-7 (Lagkouvardos et al., 2019) or *Lachnospiraceae* were dramatically reduced in *Flg*^*-/-*^ versus control mice. In addition, we observed significant reductions of few specific species of the *Ruminococcaceae, Lactobacillaceae and Rikenellaceae families*. Compared to these results, the alterations of skin microbiome observed in *Flg*^*ft/ft*^ BALB/c congenics were clearly more profound with a striking over-representation of *Firmicutes*.

Collectively, we find that loss of Flg in mice results in mild immune activation in the skin that increases with age and reduced diversity of skin microbiota.

### BALB/c *Flg*^*-/-*^ skin shows slight increase in T cell compartment but otherwise unchanged immune cell populations

To assess effects of the loss of filaggrin on the local immune system, we quantified immune cells in BALB/c *Flg*^*-/-*^ skin by flow cytometric analysis of skin cell suspensions. Importantly, CD45^+^ hematopoietic cells were not increased in relation to non-hematopoietic skin cells (mostly keratinocytes), confirming the absence of inflammatory infiltration observed histologically (Fig. 4A). While *Flg*^ft/ft^ BALB/c congenics feature larger eosinophil, mast cell and ILC2 populations (Saunders et al., 2016), we did not observe any increase in eosinophils, mast cells, neutrophils or ILC2s in BALB/c *Flg*^*-/-*^ mice versus controls (Fig. 4B and E). We also found CD86 expression by CD11c^+^ cells and the size of the intraepidermal Langerhans cell population unchanged in BALB/c *Flg*^*-/-*^ compared to control animals (Fig. 4C and D). We consistently observed a moderate increase in T cells that was most likely accounted for by expansion of the γδ T cell population (Fig. 4F).

**Figure 4.**
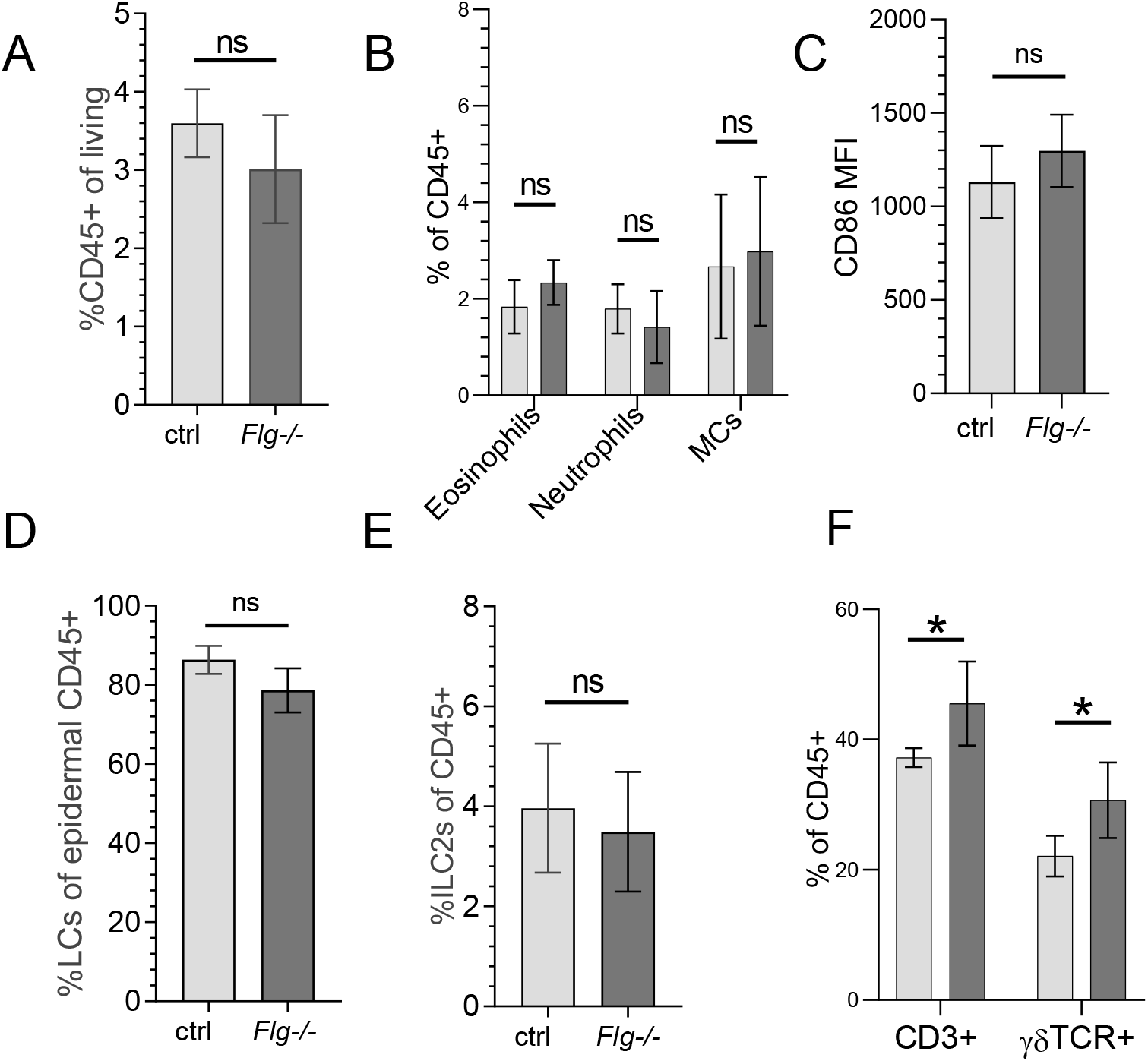
Flow cytometric analysis of immune cells in skin cell suspensions from 8-12 wk-old BALB/c *Flg*^*-/-*^and control female mice. N=5-6 both groups. A) Fraction of CD45^+^ cells in viable ear skin cells. (Cell suspension was generated by digestion of total minced ears. Absolute numbers of CD45-negative cells, mostly keratinocytes were not significantly different between mutant and control skin, not shown). Means ± SD. B) Fractions of CD11b^+^SiglecF^+^ eosinophils, CD11b^+^Gr-1^hi^ neutrophils and cKit^+^FceRI^+^ mast cells (MCs) among CD45^+^ cells. Means ± SD. C) Mean CD86 expression on CD45^+^CD11c^+^ cells. Means ± SD. D) Fraction of EpCAM^+^CD45^+^ Langerhans cells among total viable epidermal cells. Cell suspension was generated by digestion of epidermal sheets. Means ± SD. E) Fraction of CD3^-^Thy1^+^IL7Ra^+^ICOS^+^CD25^+^ ILC2s among CD45^+^ cells. Means ± SD. F) Fractions of total CD3^+^ T cells and γδ T cells among CD45^+^ cells. Means ± SD, *p< 0.05.

In summary, Flg-deficient mice on pure BALB/c background do not feature increased numbers of total immune cells in the skin. While we detected some increase in skin T cell numbers, these mice do not display the key inflammatory changes reported for young *Flg*^*ft/ft*^ BALB/c congenics, in particular the accumulations of ILC2s, eosinophils and neutrophils (Saunders et al., 2016).

### No sign of atopy in BALB/c *Flg*^*-/-*^ mice

BALB/c *Flg*^*-/-*^ mice developed the ichthyotic phenotype with scaly tail and hyperlinearity, but did not show overt skin inflammation at any time point (Fig 2A and S2). To test the *Flg* mutants for systemic atopy, total serum IgE concentrations were quantified by ELISA. In contrast to *Flg*^ft/ft^ BALB/c congenics (Saunders et al., 2016), IgE levels of our BALB/c *Flg*^*-/-*^ mice at 8, 12 and 25 weeks of age did not differ significantly from levels in BALB/c controls (Fig. 5A). As the skin microbiome is an important environmental trigger factor for AD with important roles for *S. aureus* (reviewed in (Geoghegan et al., 2018)), we tested whether colonization of Flg-deficient mice with *S*.*aureus* isolates from AD patients would result in skin inflammation and systemic atopy. *Flg*^*-/-*^ animals were painted on the shaved trunk (5 times at two day intervals) with a suspension of a *S*.*aureus* CC1 isolated from an AD patient. The mice were monitored for 7 weeks thereafter, however, no overt skin inflammation (not shown) or increase in total serum IgE (Fig. 5B) was observed. As skin colonization can be impeded by the resident flora, we repeated the *S*.*aureus* colonization with prior systemic antibiotic treatment, but also this regime did not trigger skin inflammation or enhanced IgE production in our BALB/c *Flg*^*-/-*^ mice (Fig. 5B). To test whether the altered cutaneous immune milieu of *Flg*^*-/-*^ mice (Fig. 3A-D) would drive abnormal IgE responses upon exposure to exogenous antigen through the skin, we epicutaneously immunized BALB/c *Flg*^*-/-*^ mice with OVA. OVA-specific IgE was not significantly different between mutant and BALB/c control littermates 7 weeks later (Fig. 5C). We next repeated the epicutaneous OVA immunization in *Flg*^*-/-*^ or control BALB/c that had been transferred OVA-specific DO11.10 transgenic (Murphy et al., 1990) IL-4eGFP (4get) T reporter cells (Mohrs et al., 2001). As our finding that TEWL increased in older (16-18 weeks old), but not in younger mice (8-10 weeks old), indicated that the barrier defect develops with age, we included mice of both age groups. Application of OVA to the skin of DO11.10 T cell recipients did not cause any skin alteration in wild-type, or younger *Flg*^*-/-*^ recipients (not shown), whereas marked skin inflammation was elicited in 16-week-old *Flg*-deficient recipient mice (Fig. 5D). To further investigate the differences in the development of skin inflammation in DO11.10 cell recipient *Flg*-deficient but not wild-type recipient mice, we assessed DO11.10 cellular responses in the skin draining LNs (Fig 5.E). In the LNs of Flg-deficient mice, but not control recipient wild-type mice, there was vigorous cell proliferation with elevated frequencies of IL-4-eGFP^+^ cells) indicating enhanced OVA-specific Th2 cellular response in skin LN of *Flg*^*-/-*^ (Fig. 5E).

**Figure 5.**
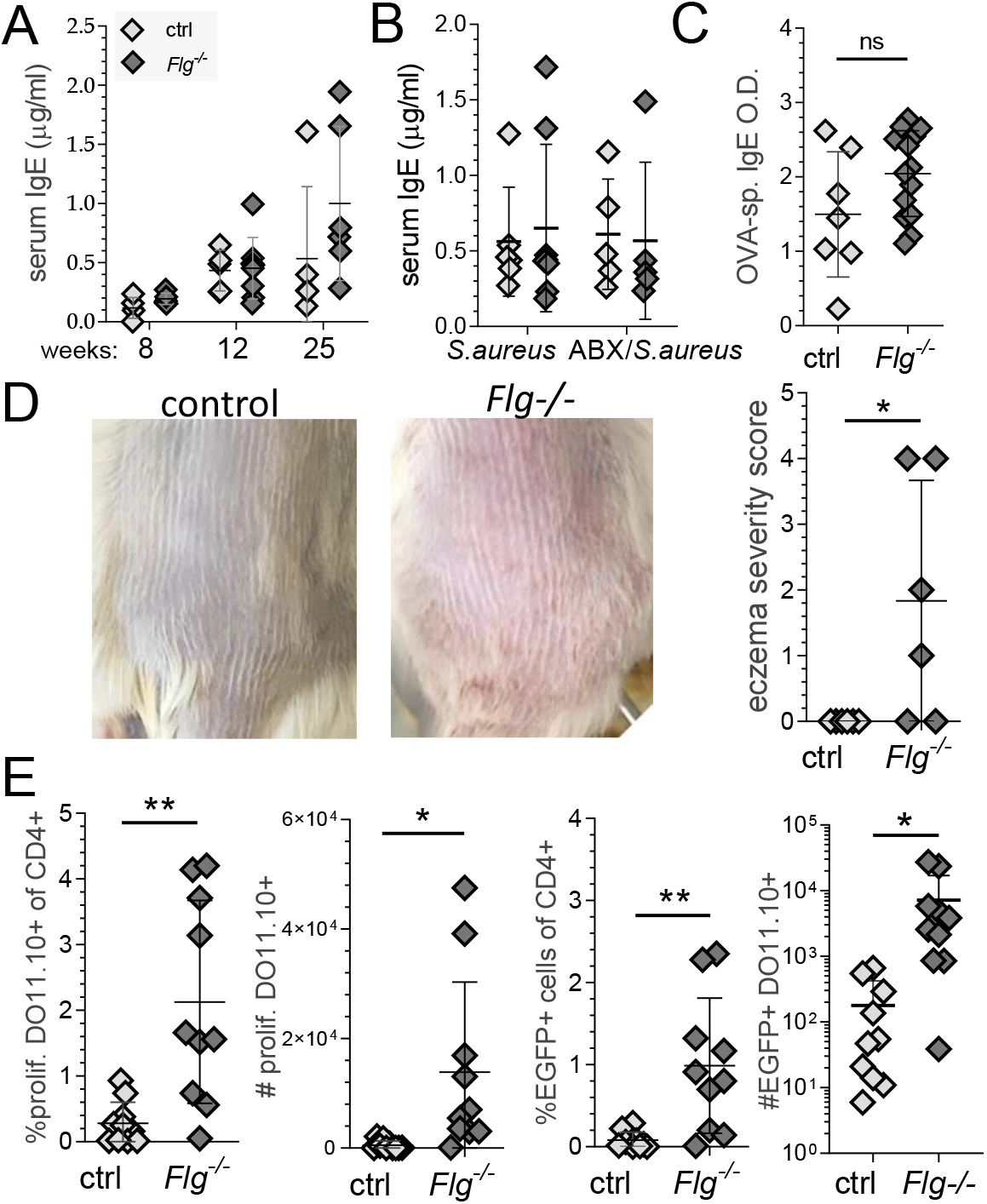
Loss of Flg in mice of pure BALB/c background does not result in atopy A) Quantification of total serum IgE of BALB/c *Flg*^*-/-*^(n=5-8) and littermate control mice (n=5) by ELISA at indicated age. Means ± SD. B) Serum IgE levels (as in A) of 14-16 week-old BALB/c *Flg*^*-/-*^ (n=5-8) and littermate control mice (n=5 or 6) colonized with a *S*.*aureus* isolate from an AD patient (CC1 strain) without or with prior systemic antibiotic treatment (ABX). *S*.*aureus wt group n=6*, ko group n=8. ABX/ *S*.*aureus* groups n=5 both genotypes. Means ± SD. C) Quantification of OVA-specific serum IgE levels (ELISA) of 16 week-old BALB/c *Flg*^*-/-*^(n=14) and littermate control mice (n=7) immunized epicutaneously with OVA. Means ± SD. D and E) Immune response of BALB/c *Flg*^*-/-*^recipients of OVA-specific transgenic DO11.10 T cells to epicutaneous OVA immunization. 16-week-old mice received i.v. transfer of 2×10^6^ MACS-purified, splenic CD4^+^ T cells isolated from transgenic DO11.10 mice, stained with proliferation dye (eFluor™ 670) on day 0 and were epicutaneously treated with OVA on shaved back on days 1, 2 and 3. D) Macroscopic assessment of skin inflammation on day 5. Left: representative images of back skin, right: dermatitis score (see Fig. S5), n=5 for controls. Means ± SD, * p<0.05. E) Proliferative response and IL-4 expression of donor DO11.10 cells. Frequency (left) of proliferating transgenic cells (see Fig. S5 for gating) in total CD4^+^ T cells, (middle left) total number of proliferating transgenic cells in the draining LNs and (middle right) and frequency of IL-4 transcriptional reporter (4get) expressing cells among CD4^+^ T cells of the draining LN and their absolute numbers (right) (see Fig. S5 for gating). N=5 mice both groups with 2 LNs analysed per mouse, means ± SD, * p<0.05, ** p<0.01. Data representative of two experiments.

In summary, we show that loss of Flg in mice of pure BALB/c background does not result in increased total serum IgE, neither spontaneously nor after colonization with *S*.*aureus*. Also epicutaneous immunization did not elicit more specific IgE in BALB/c *Flg*^*-/-*^ mice compared to control mice, but did induce enhanced T cell proliferation and skin inflammation in recipients of antigen-specific transgenic T cells.

### Accurate whole genome sequencing reveals that *Flg*^*ft/ft*^ BALB/c congenic mice are in fact *Flg*^*ft/ft*^ *Tmem79*^*ma/ma*^ double mutants, explaining their atopic phenotype

Our finding that complete loss of Flg in mice of pure BALB/c background did not result in detectable atopy, contrasting with the robust atopic phenotype of *Flg*^*ft/ft*^ BALB/c congenics (Saunders et al., 2016), suggested that further genetic variation within the congenic interval in addition to the *Flg* loss-of-function mutation is essential for development of atopy in FLG-deficient animals.

We therefore generated a highly accurate whole genome sequence of the *Flg*^*ft/ft*^ BALB/c congenic strain and our newly generated BALB/c *Flg*^*-/-*^ mice using PacBio long read single-molecule real-time sequencing (Eid et al., 2009; Wenger et al., 2019)). Average high-fidelity (HiFi) long read coverage was 16.7x for the *Flg*^*ft/ft*^ BALB/c congenic strain and 14.7x for the BALB/c *Flg*^*-/-*^ strain. Both strains were assembled using the *de novo* assembler hifiasm (version 0.7). We used the BALB/c *Flg*^*-/-*^assembly to map the read sets of both genomes for comparison of variations. To identify chromosome 3 contigs, the BALB/c *Flg*^*-/-*^ assembly was mapped to the GRCm38.p6 reference with minimap2 (version 2.17-r941). Annotation was performed by lift-over from the M. musculus GRCm38 genome built 6. Chromosome 3 alignments were used for variant analysis followed by variant effect prediction. Both *de novo* assemblies of chromosome 3 and annotations can be downloaded from https://bds.mpi-cbg.de/hillerlab/Bat1KPilotProject/.

In the *Flg*^*ft/ft*^ BALB/c congenic sequence, we identified the Flg 5303delA frame shift mutation in repeat 6, as expected, and the sequence of the new BALB/c *Flg*^*-/-*^ line revealed the size of the deletion we had introduced by CRISPR/Cas9-mediated mutagenesis (Fig. 1). Alignment of the new chromosome 3 sequence of *Flg*^*ft/ft*^ BALB/c congenic and BALB/c *Flg*^*-/-*^ mice demonstrated that chromosome 3 (and all other chromosomes, not shown) of the two strains were nearly identical with our BALB/c *Flg*^*-/-*^ de novo assembly, with an average of 1 variation per 9694 bp outside of the congenic interval (QV score 39.9). However, the two sequences extensively differed within a 51.3 MB congenic interval, with an average of 1 variation per 145 bp (QV score 21.6) (Fig. 6). This interval was in roughly equal parts derived from C3H (27 MB, near perfect match with C3H in all positions for which published C3H sequence information was available) and from C57BL/6 (24.3 MB) in the *Flg*^*ft/ft*^ BALB/c congenic strain. These findings reflect the history of sequential backcrossing of the *Flg*^*ft/ft*^ mutation first to C57BL/6 (Saunders et al., 2013) and then to BALB/c (Saunders et al., 2016). Within the 51.3 MB congenic interval we identified 610 genes having functional annotations. Of these, 12 differed between the *Flg*^*ft/ft*^ BALB/c congenic and the BALB/c *Flg*^*-/-*^ strain by deleterious mutations (frame shift, gain or loss of stop codon, loss of start codon or loss of splice site), while 143 genes differed by amino acid exchange mutations predicted to potentially impact on structure or function (Table S1).

**Figure 6.**
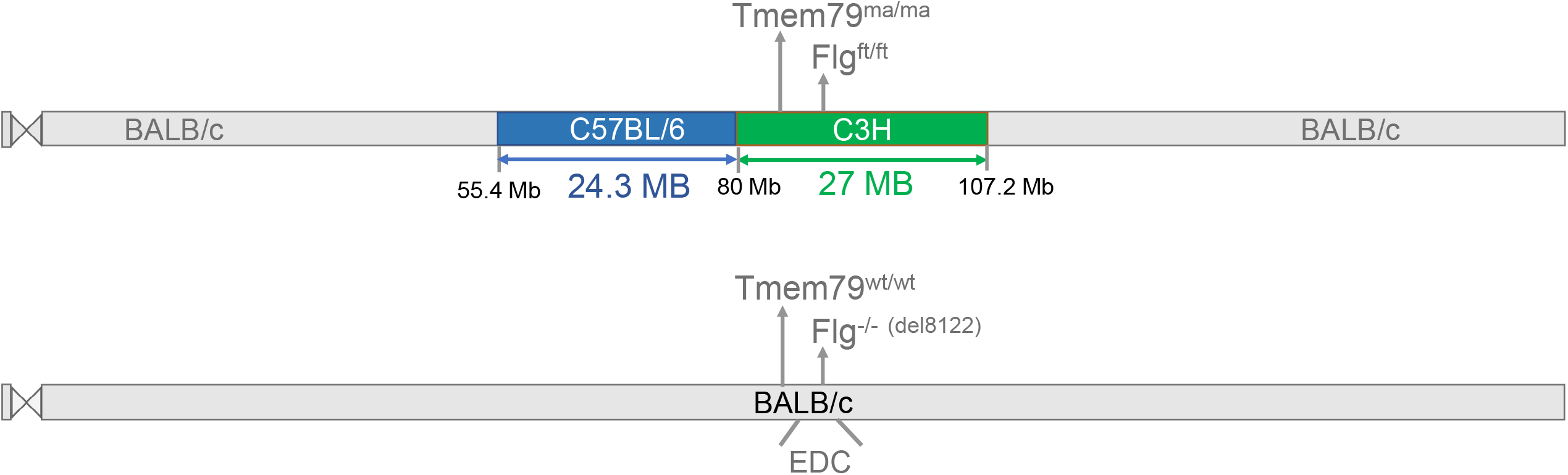
Unexpected presence of the *Tmem79*^*ma*^ gene variant in the *Flg*^*ft/ft*^ BALB/c congenic strain An accurate sequence of BALB/c chromosome 3 was generated by whole genome long read single molecule real-time sequencing (PacBio) of our new BALB/c *Flg*^*-/-*^ strain and de novo assembly. In parallel, the *Flg*^*ft/ft*^ BALB/c congenic strain (Saunders et al., 2016) was sequenced and chromosome 3 sequences of the two strains were compared. The two strains differ by a congenic interval of approximately 51.3 MB that is made up by a C57BL/6- and a C3H-derived region, the latter containing *Flg* and *Tmem79*. As expected, the *Flg* gene of the *Flg*^*ft/ft*^ BALB/c congenic strain (Saunders et al., 2016) harbors the *Flg*^*ft*^ mutation (1bp deletion in *Flg* repeat 6) while our BALB/c *Flg*^*-/-*^strain carries an 8122 bp (out-of-frame) deletion in *Flg* exon 3. Unexpectedly, the *Flg*^*ft/ft*^ BALB/c congenic strain (Saunders et al., 2016) is homozygous for the atopy-associated *Tmem79*^*ma*^ mutation that was reported to be crossed out of this strain before initiation of backcrossing to BALB/c (Saunders et al., 2013). EDC, epidermal differentiation cluster.

Among the deleterious mutations, we unexpectedly found, the Y280Stop mutation of the *Tmem79/mattrin* gene (Saunders et al., 2013), located in the CH3-derived interval of the *Flg*^*ft/ft*^ BALB/c congenic mice. The *Flg*^*ft*^ mutation originally arose on a strain (mixed background, partially C3H) with matted hair phenotype to yield the flaky tail mouse (Fallon et al., 2009; Lane, 1972; Presland et al., 2000). Saunders et al. identified the Y280Stop mutation of *Tmem79*, a gene in linkage disequilibrium with *Flg* on chromosome 3, to be responsible for the matted phenotype and re-named the gene *mattrin* (Saunders et al., 2013). In order to separate the *Flg*^*ft*^ and the *Tmem79*^*ma*^ mutations, Saunders et al. bred flaky tail mice to C57BL/6 and reported that homozygous *Tmem79*^*ma/ma*^ mice (wild type for Flg) showed severe systemic atopy with eczema, spontaneous increase in IgE levels and asthma (Saunders et al., 2020; Saunders et al., 2013), and also identified a human *TMEM79* missense SNP associated with human AD (Saunders et al., 2013). *Flg*^*ft/ft*^ C57BL/6 mice (assumed to be wild type for *Tmem79*) were backcrossed extensively to BALB/c to yield the *Flg*^*ft/ft*^ BALB/c congenic strain that is characterized by a prominent atopic phenotype, including eczema, high IgE and spontaneous asthma (Saunders et al., 2016; Schwartz et al., 2019). Our sequencing result, however, demonstrates that the *Flg*^*ft/ft*^ BALB/c congenic strain is homozygous for the *Tmem79*^*ma*^ mutation, indicating that either the two mutations had not been successfully separated, or that the *Tmem79*^*ma*^*Flg*^*ft*^ haplotype was accidentally reintroduced during backcrossing to BALB/c.

The phenotype of mice with isolated lack of Tmem79 is strikingly similar to that of the *Flg*^*ft/ft*^ BALB/c congenics. Young adult *Tmem79*^*ma/ma*^ mice (Saunders et al., 2020) develop eczema with the same kinetics, macroscopic appearance and severity as reported for the *Flg*^*ft/ft*^ BALB/c congenics. Neonatal ichthyosis does not occur in *Tmem79*^*ma/ma*^ mice due to intact Flg. *Tmem79*^*ma/ma*^ mice also reproduce the high serum IgE and spontaneous lung inflammation of *Flg*^*ft/ft*^ BALB/c congenics (Saunders et al., 2020). Genotyping of *Flg*^*ft/ft*^ BALB/c congenic mice for the *Tmem79*^*ma*^ and *Flg*^*ft*^ mutations confirmed the sequencing data, with both *Tmem79* and *Flg* mutations detected. It is therefore, indicative that the atopic phenotype of *Flg*^*ft/ft*^ BALB/c congenics is caused by the *Tmem79*^*ma*^ mutation. However, also differences in other genes located in the congenic interval could contribute to and modulate skin inflammation and atopy in *Flg*^*ft/ft*^ BALB/c congenic mice. The list of genes for which the *Flg*^*ft/ft*^ BALB/c congenics and our BALB/c *Flg*^*-/-*^ animals show relevant (deleterious or structurally/functionally damaging) differences contains several genes with functions in skin barrier, including *crnn* (encoding cornulin), *Flg2*, and *Ivl* (endoding involucrin), and genes with functions in the immune system, such as *Fcgr1, Pglyrp3* and *Arnt* (Table S1)

Collectively, we show that the decisive difference between non-atopic BALB/c *Flg*^*-/-*^mice and the atopic *Flg*^*ft/ft*^ BALB/c congenic strain is the unexpected presence of the atopy-causing *Tmem79*^*ma*^ mutation in the latter.

## Discussion

To resolve the unexplained discrepancy between prominently atopic Flg-deficient *Flg*^*ft*^ BALB/c congenic mice (Saunders et al., 2016) and mice with targeted inactivation of Flg (Kawasaki et al., 2012), and to determine the effects of Flg deficiency on skin immune regulation, we generated Flg-deficient mice on a pure BALB/c background by inactivating the Flg gene in BALB/c embryos. These animals do not develop systemic atopy. We sequenced the genome of the atopic *Flg*^*ft*^ BALB/c congenics (Saunders et al., 2016) and discovered that these animals unexpectedly harbor the atopy-causing matted mutation of the *Tmem79* gene (*Tmem79*^*ma*^) that is linked to the *Flg* locus. The genome sequencing data was validated by genotyping, with both *Tmem79*^*ma*^ and *Flg*^*ft*^ mutations present in the *Flg*^*ftft*^ BALB/c congenic mice. These findings shed light on the hitherto puzzling and conflicting published results on skin barrier-defective mouse models.

In accordance with the phenotype of *Flg* knock out mice (Kawasaki et al., 2012), our BALB/c *Flg*^*-/-*^mice demonstrate that, in mice, loss of Flg results in dry and scaly skin only postnatally, while skin and fur are macroscopically normal throughout adult life. Histologically, substantial reduction of keratohyalin granules is the only prominent change. Loss of Flg alone does not result in spontaneous development of overt inflammation as our BALB/c *Flg*^*-/-*^ mice, like the published *Flg* knock out line (Kawasaki et al., 2012), show no macroscopic dermatitis. Also histologically, we could not detect any inflammatory skin changes in line with Kawasaki et al., who reported no inflammatory infiltration (Kawasaki et al., 2012). Flg-deficiency, however, is clearly not immunologically inert. As was reported in the published *Flg* knock out line (Kawasaki et al., 2012), we observed enhanced adaptive responses to epicutaneously administered antigen, most likely reflecting abnormal antigen penetration. We also found spontaneous changes in the skin immune system. Slight alterations in BALB/c *Flg*^*-/-*^ keratinocyte gene expression suggest local immune activation. Moreover, while total numbers of immune (CD45^+^) cells in the skin were not increased in skin of Flg-deficient mice, we observed, changes in numbers of skin-resident immune cells by flow cytometry, i.e. a moderate expansion of T cells. However, the marked increases in ILC2s, eosinophils and mast cells that characterizes the *Tmem79*^*ma/ma*^*Flg*^*ft/ft*^ BALB/c congenics (Saunders et al., 2016; Schwartz et al., 2019) do not occur in our BALB/c *Flg*^*-/-*^ mice. Absence of Flg results in no detectable change of TEWL in young adults, however, we found water loss increased in older mice. To test whether increased TEWL in older animals correlated with increased Ag penetration, we epicutaneously immunized 8 week-old (not shown) and 16 week-old Flg-deficient and control mice. No sign of skin inflammation was observed in control mice or in 8-week-old flg-deficient mice. However, in 16-week-old *Flg*^*-/-*^mice we observed eczematous skin inflammation, increased OVA-specific T cell proliferation and increased IL-4 production in the skin draining lymph nodes. In addition, we found a sponantous induction of *IL-1b* transcription in total skin, suggesting that age related loss of epidermal barrier resulted in immune activation. The microbiome of BALB/c *Flg*^*-/-*^skin clearly differs from microbiomes of control skin with reduced diversity and systematic reduction of particular species. However, these changes are less pronounced and qualitatively different compared to the gross skin microbiome alteration in *Tmem79*^*ma/ma*^*Flg*^*ft/ft*^ BALB/c congenics (Saunders et al., 2016; Schwartz et al., 2019) which display a striking overrepresentation of firmicutes (Schwartz et al., 2019). The effects of filaggrin deficiency on skin microbiota may either be a direct consequence of the lack of Flg and resulting changes of skin pH, moisture and turnover or texture of the cornified layer, or could be a result of the subtle immune-activation we observed in the BALB/c *Flg*^*-/-*^ animals. Obviously, additional loss of Tmem79 will also significantly impact the skin microbiome.

Importantly, we now show that the atopic phenotype of *Flg*^*ft/ft*^ BALB/c congenics (Saunders et al., 2016) with dermatitis, increased serum IgE and spontaneous asthma, is likely caused by the *Tmem79*^*ma/ma*^ mutation that is linked to the *Flg*^*ft*^ allele and was not known to be present in this strain. Mice with isolated Tmem79 deficiency develop a largely similar phenotype (Saunders et al., 2020; Saunders et al., 2013), which is in sharp and previously unexplained contrast to the complete lack of atopy in the published *Flg* knock out line (Kawasaki et al., 2012) and our BALB/c *Flg*^*-/-*^ animals. Both lines feature neither dermatitis, increase in serum IgE, nor lung inflammation. Not even after colonization with *S*.*aureus* isolates from AD patients did IgE levels rise in BALB/c *Flg*^*-/-*^animals.

Future experiments are required to elucidate whether lack of *Flg*, while not causing atopy alone, nevertheless modulates skin inflammation and systemic atopy of Tmem79-deficient mice. Comparison of findings in *Tmem79*^*ma/ma*^*Flg*^*ft/ft*^ BALB/c congenic mice (Saunders et al., 2016; Schwartz et al., 2019) and in *Tmem79*^*ma/ma*^ C57BL/6J congenic animals (Saunders et al., 2020; Saunders et al., 2013) are confounded by the difference in strain background. The difference in skin phenotype between the mice could be caused by the absence of Flg. Alternatively, however, additional genetic variation in the respective congenic interval could exert modulating effects on skin inflammation and atopy of *Tmem79*^*ma/ma*^*Flg*^*ft/ft*^ mice. It is noteworthy that the genes significantly altered in *Tmem79*^*ma/ma*^*Flg*^*ft/ft*^ mice include several barrier genes (Table S1), but also genes with known functions in the immune system.

Defects of the epidermal skin barrier one of the most important predisposing factor for atopic disease, notably not only for atopic dermatitis, but also for systemic atopy, i.e. allergic sensitizations and asthma (Palmer et al., 2006). FLG deficiency, the cause of ichthyosis vulgaris, is arguably the most important barrier defect associated with atopy. Our finding that lack of Flg is not sufficient to cause eczema and systemic atopy in mice is line with the fact that a large fraction of IV patients homozygous for *FLG* null alleles never develop atopic dermatitis, allergic sensitizations or asthma. Our result highlights the essential role of atopy promoting variants of other genes required to synergize with FLG deficiency in causing the atopic predisposition. Also in human atopy, defects of other ‘barrier genes’ linked to FLG in the EDC and adjacent regions may contribute to the atopic phenotype. Mouse models with defects of one or several ‘skin barrier genes’, including the *Tmem79*^*ma/ma*^*Flg*^*ft/ft*^ BALB/c congenics, will be instrumental in understanding systemic immune dysregulation in atopy associated with compromised skin barrier.

## Acknowledgments

This work was funded by DFG grants RO2133/9-1 to A.R. and DA 1311/3-1 to A.D. in the setting of FOR2599 and was supported by Federal Ministry of Education and Research (BMBF grant 01IS18026C). We thank Werner Müller for helpful discussion. Christina Hiller, Livia Schulze, Madelaine Rickauer and Tobias Häring provided expert technical assistance.

## Legends

**Figure S1.**
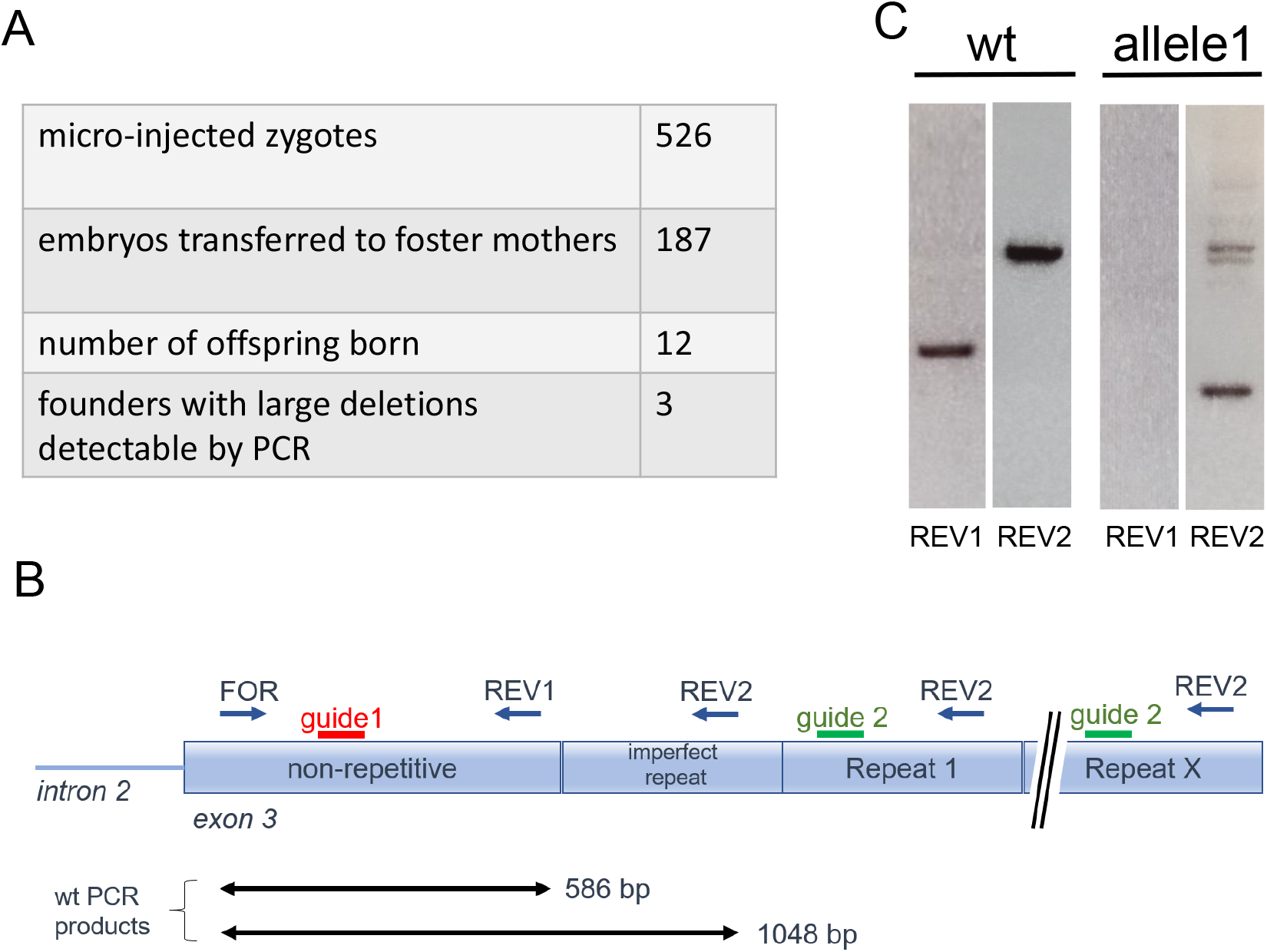
Generation and genotyping of BALB/c *Flg*^*-/-*^ mice A) Numbers of BALB/c zygotes injected with Cas9-RNPs, transferred embryos, number of viable offspring resulting from these transfers and numbers of founders harboring large deletions involving cleavage by both guides. B) Strategy for PCR-based identification of mutant *Flg* alleles. Separate reactions were performed with primer pairs FOR-REV1 and FOR-REV2. C) PCR result for mutant allele 1, resulting from deletion downstream of the guide 1 target site and one of the guide 2 target sites with loss of *Flg* repeats. The deletion results in absence of the FOR-REV1 product and alters the length of product FOR-REV2.

**Figure S2.**
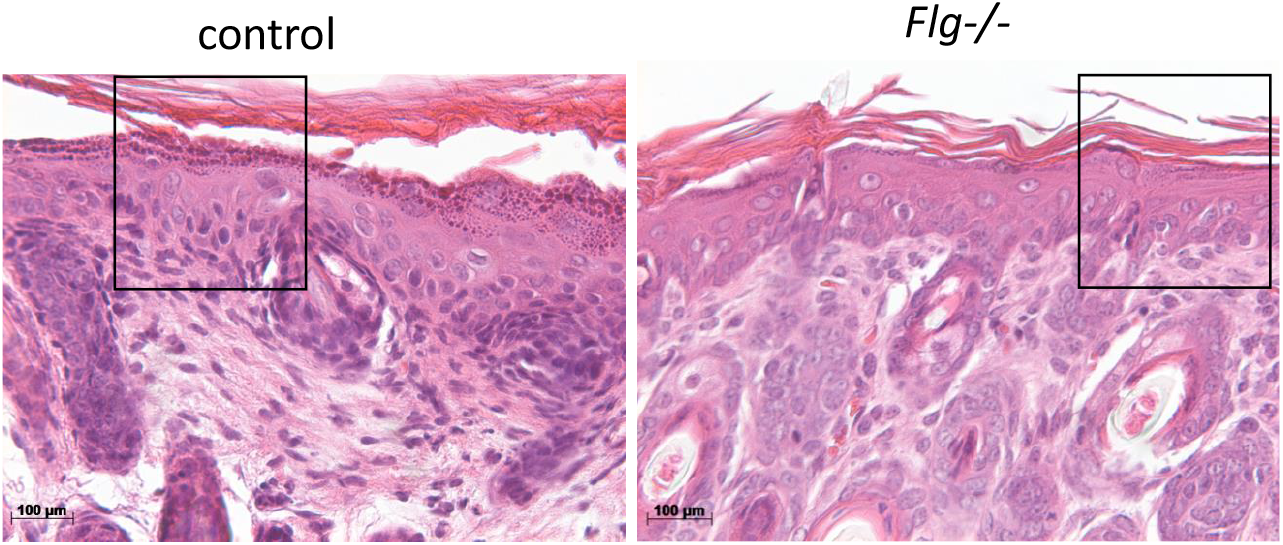
Ear skin histology of BALB/c *Flg*^*-/-*^ and control mice. H&E staining. Insets indicate the images shown in Fig. 2A. Images representative of at least 10 mice. Scale bar 100 µM.

**Figure S3.**
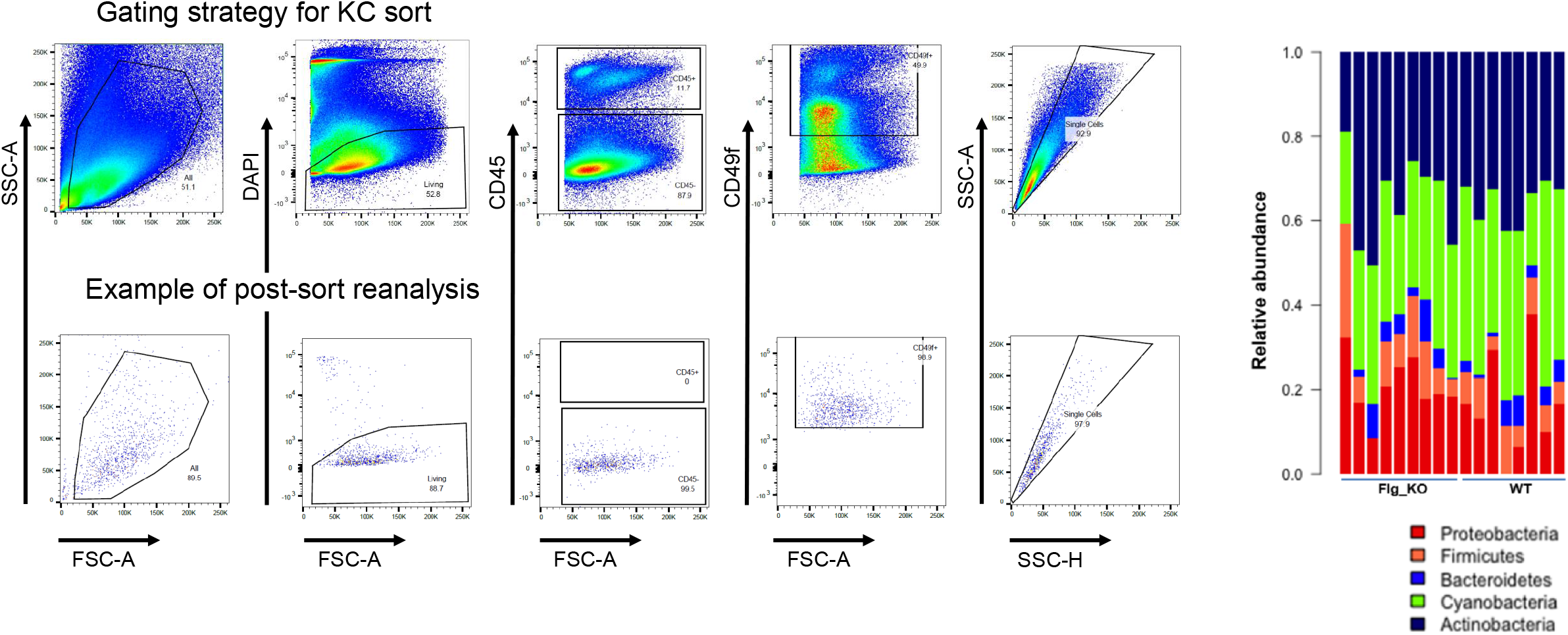
A) Gating strategy and post-sort reanalysis of the flow cytometric purification of basal (CD49f^+^) keratinocytes. B) Relative abundance of microbial phyla in cheek skin swabs from 8 week-old female BALB/c *Flg*^*-/-*^ and control co-housed females

**Figure S4.**
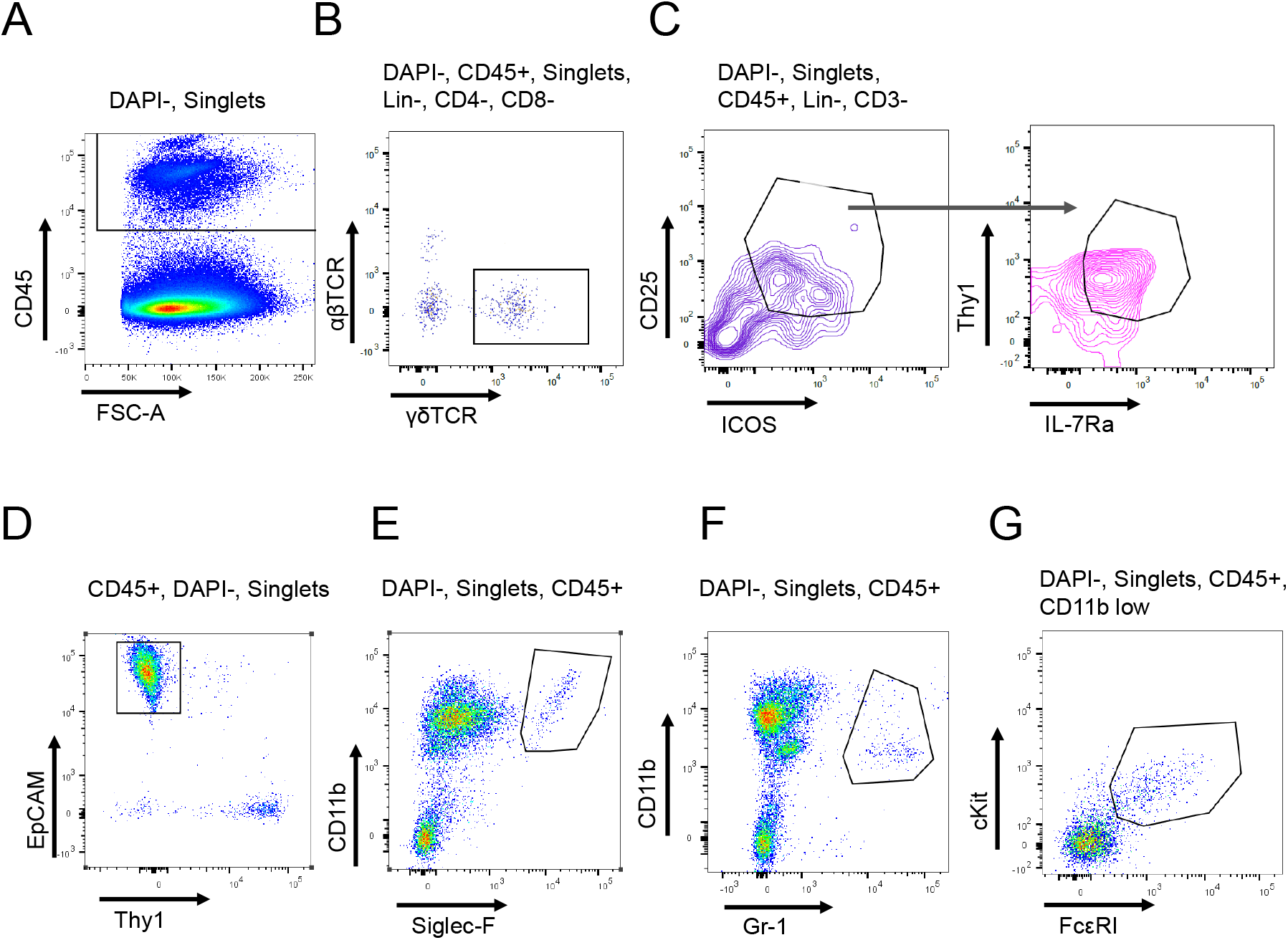
Gating strategies for quantification of individual immune cell populations in total skin cell suspensions or epidermal cell suspensions (in the case of Langerhans cells). A) Whole ear skin CD45^+^ cells, B) γδTCR^+^ T cells, C) ILC2 population, D) epidermal Langerhans cells, E) eosinophils, F) neutrophils, G) mast cells.

**Figure S5.**
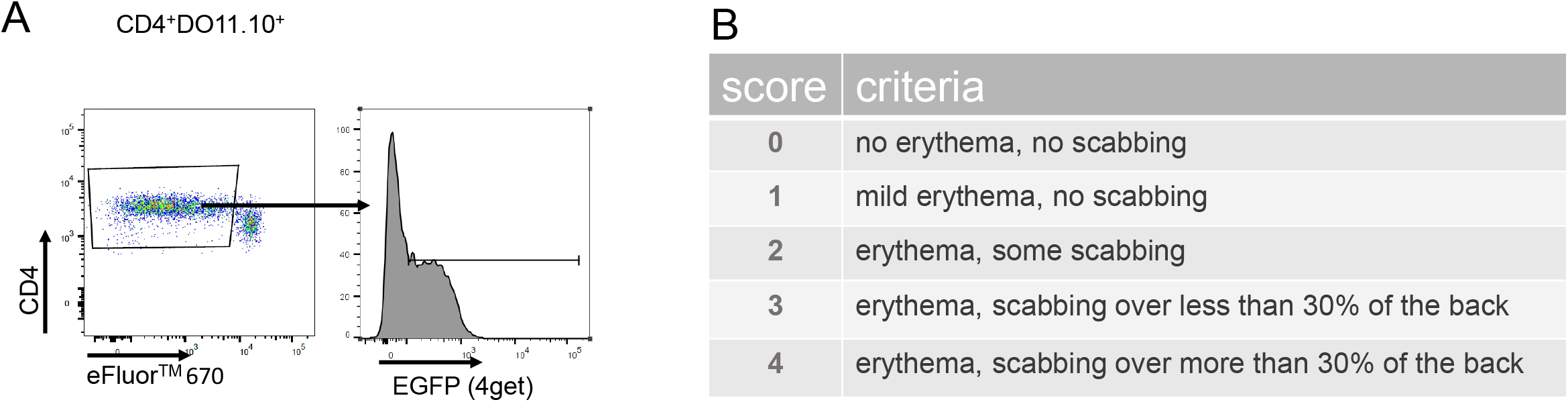
A) Gating of proliferating and IL-4-expressing OVA-specific transgenic DO11.10 donor T cells in skin draining lymph node upon epicutaneous OVA administration. B) Eczema severity score used to macroscopically quantify dermatitis in BALB/c *Flg*^*-/-*^ and control recipients of OVA-specific transgenic DO11.10 T cells epicutaneously treated with OVA (main Figure 5D).

**Table S1.**
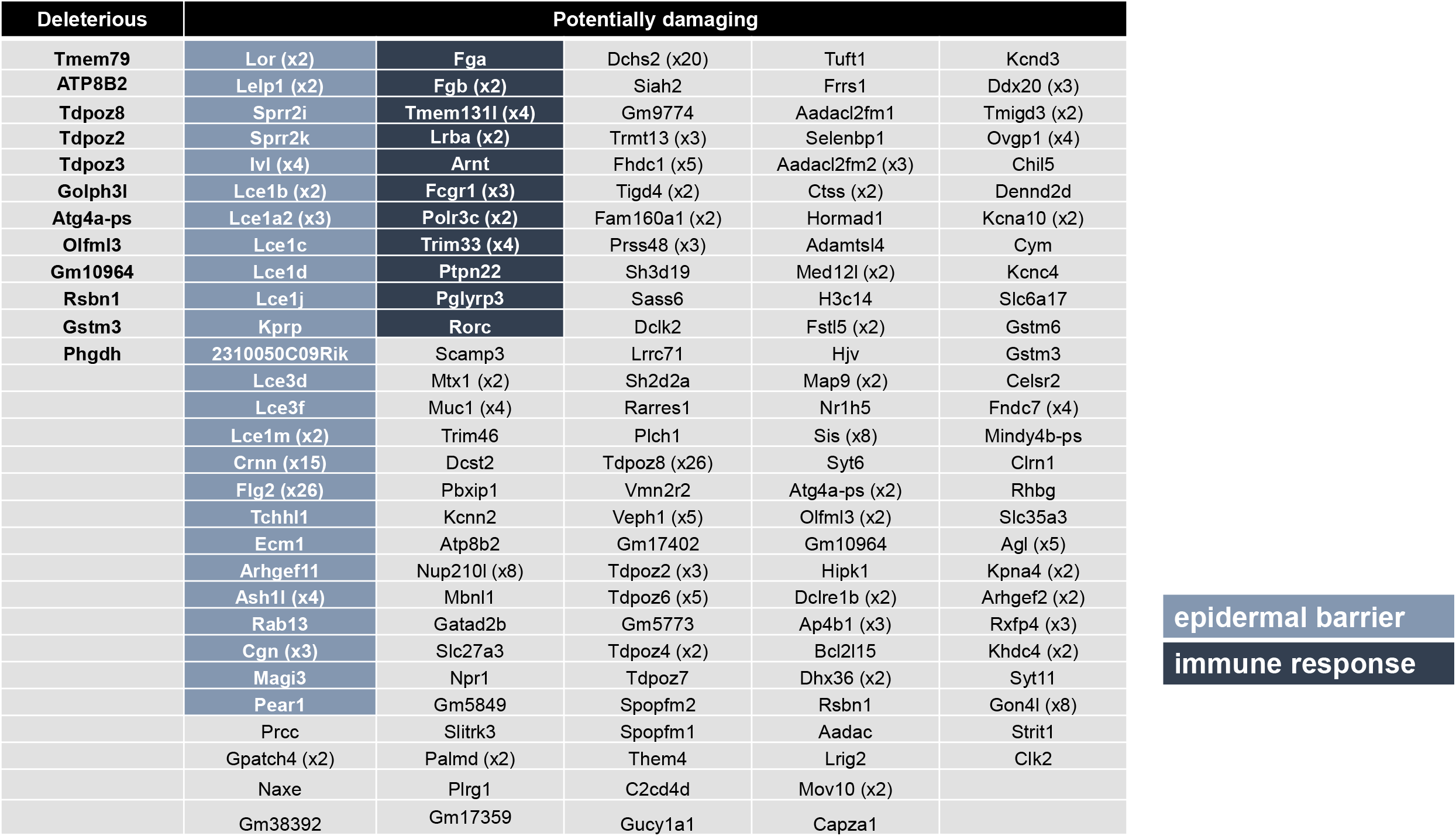
Genes located in the congenic interval of *Flg*^*ft/ft*^ BALB/c congenic mice showing deleterious or structurally/functionally relevant differences between *Flg*^*ft/ft*^ BALB/c congenics and BALB/c mice. ‘Deleterious’: gain or loss of Stop codon, loss of a Start codon, frame shift, loss of splice site; ‘potentially damaging’: damaging missense mutations as predicted by SIFT. Numbers in brackets indicate numbers of individual potentially damaging mutations present in respective genes.

## Material and methods

### Mice

Animals were kept under specific pathogen-free conditions (SPF) in individually ventilated cages (IVCs) at the Experimental Centre, Medical Faculty Carl-Gustav Carus, Technische Universität Dresden. Pathogen-free conditions were regularly tested according to the Federation for Laboratory Animal Science (FELASA). All animal experiments were done according to the German animal welfare law approved by the Landesdirektion Dresden (ref. no. DD25-5131/474/24) Wt BALB/c mice were obtained from Janvier. *Flg*^*ft/ft*^ BALB/c congenic mice were kept at Trinity College Dublin and genotype for mutations in *Flg*^*ft*^ (5303delA) and *Tmem79*^*ma*^ (p.Y280*) as described Saunders 2013 (Saunders et al., 2016).

### Generation and genotyping of *Flg*^*-/-*^ mice on a pure BALB/c background

Zygotes were isolated from superovulated female BALB/c mice at E0.5. 526 single zygotes were microinjected with Cas9 RNPs prepared from 5,25 µM NLS-Cas9 (Toolgen), 1,73 µM crRNA1 (unique target site: 5’ GCTGGCAAAAGCATATTATG-3’), 1,73 µM crRNA2 (target site in each of the *Flg* repeat regions: 5’ GTCAGCGCAAGATCAGGCTC-3’) and 6,9 µM tracrRNA (all IDT). 187 embryos developed to the two-cell stage and were transferred into pseudopregnant foster mothers. Genotyping of *Flg*^*-/-*^ mice was performed by PCR. Primers were designed to bind the BALB/c *Flg* gene in the non-repetitive region of exon 3 (For *5’CCAGATGACCAAGACATCGCTG 3’* and Rev1 *5’CTCTGGGTCTTCTGTTTCCCTCTC 3’*) or the *Flg* repeat regions (Rev2 *5’ GGCCCTGTGCTTGGCCTTG 3’*) (Fig S1).

### Western blot analysis

Ears and tail skin were collected from 8 week-old mice and minced. Protein extraction was performed by 1h incubation at 95°C in lysis buffer (6% SDS, 30% glycerol, 10mM DTT, bromophenol blue) agitation. Upon SDS-PAGE, transfer to nitrocellulose membrane (Amersham™ Protran® Premium 0.45 µm NC) and blocking (5% BSA/TBST, 2h at RT), membranes were incubated with primary antibodies for 1h at RT with shaking (1:2000 anti-filaggrin clone Poly19058, Biolegend; 1:2000 anti-β-actin, clone 13E5, Cell Signaling Technology). After washing (3×5 min in Tris-buffered saline containing 0.1% v/v Tween20, with shaking), primary antibody was detected with alkaline phosphatase–conjugated polyclonal goat anti-rabbit IgG, D0487, Dako Denmark A/S) and alkaline phosphatase substrate (NBT/BCIP, Roche).

### Filaggrin immunofluorescence staining

Tail skin was fixed in 4% PFA, paraffin-embedded and cut into 5 µM sections. Slides were heat treated for 20 minutes at 701C, followed by deparaffinization and rehydration. Sections were permeabilized by 0.1% Triton X-100/PBS treatment (10 min, RT), followed by washing. All washing was by incubating the slides 3 times in PBST Buffer for 5 min. Slides were blocked in 5% goat serum/PBST, for 1h at RT, with agitation, followed by incubation for 1h at RT in a 1:1000 dilution in blocking buffer of primary Ab (purified anti-Filaggrin polyclonal Ab, clone Poly19058, Biolegend). After washing, slides were incubated for 1 h at RT in a 1:500 dilution of secondary Ab (IgG Highly Cross-Adsorbed Goat anti-Rabbit, Alexa Fluor® 488, Invitrogen™). After the final wash, slides were mounted with mounting medium (Dako Fluorescence Mounting Medium, Dako North America, Inc.) containing 10 µg/ml of DAPI. Slides were analyzed on a LSM confocal microscope Zeiss LSM880 with AIRYscan.

### Quantification of transepidermal water loss (TEWL)

Mice that had been shaved on their back 24h earlier were anesthetized (Ketamin/Xylazin) and TEWL was measured using an MDD4 device (CK electronic) according to the manufacturer’s instructions. Each value was acquired as a mean of three individual measurements.

### Sorting of basal ear skin keratinocytes for RNA extraction

Ears were minced in 1ml of DMEM/20mM Hepes buffer, and 0.5 mL of an enzyme mixture containing Liberase (25µg/ml, Liberase™ Research Grade, Sigma Aldrich), Hyaluronidase (0.5 mg/ml Sigma, 100mg Hyaluronidase from bovine testes, Sigma Aldrich), and DNaseI (200U/µl, DNase I grade II, from bovine pancreas, Sigma Aldrich) was added, followed by incubation for 1h at 371C with agitation (1300rpm). Disassociated tissue was filtered through a 100 µm mesh, washed with 5 ml of cold FACS buffer (0.5% BSA/2mM EDTA/PBS), and pelleted by centrifugation at 1200 rpm for 5min at 41⍰C. Cells were resuspended in 75µL of Fc blocking medium (5 µg/ml Purified anti-mouse CD16/32 Antibody, Biolegend in FACS buffer), and incubated for 20 minutes on ice. After centrifugation, cells were resuspended in 75µl of FACS buffer containing surface staining antibodies, and incubated on ice for 45 min. Antibodies used for the staining are anti-mouse CD45 (PerCP-Cy5.5 labelled, clone 30-F11, eBioscience), anti-mouse CD11b (FITC labelled, clone M1/70, eBioscience), anti-mouse F4/80 (APC labelled, clone BM8, eBioscience), anti-mouse CD49f (PE labelled, clone eBioGoH3, eBioscience). After washing, stained cells were resuspended in FACS buffer containing 1µg/ml of DAPI and kept on ice. Cell sorting of DAPI low/CD45-/CD49f^+^ cells was performed on a BD FACSAria™ III instrument, using an 85 µM nozzle. Conflicting events were excluded from the sort (purity sorting). Cells were sorted directly into a lysis buffer (500µl of Buffer RLT Plus containing β-mercaptoethanol, Qiagen), and stored at -80⍰C before proceeding with RNA extraction.

RNA was extracted using RNeasy® Plus Micro Kit (Qiagen) according to the manufacturer’s instructions and stored at -80°C. The quality of the isolated RNA was determined on a Bioanalyzer instrument (Agilent Technologies, Eukaryote Total RNA Pico chip). Samples with RNA integrity number higher than 7 were selected for RNA sequencing. mRNA libraries were subjected to sequencing on an Illumina®HighSeq platform yielding 30 -60 million reads per sample. Raw reads were mapped to the mouse genome (mm10) and splice-site information from Ensembl release 87 (Zerbino et al., 2018) with gsnap (version 2018–07-04; (Wu and Nacu, 2010). Uniquely mapped reads and gene annotations from Ensembl were used as input for featureCounts (version 1.6.2; (Liao et al., 2014)) to create counts per gene. RNAseq differential expression analysis was performed with the R package DESeq2 (Love et al., 2014).

### quantitative RT-PCR

Total RNA from sorted CD45-negative cells was isolated using RNeasy® Plus Micro Kit (Qiagen), while RNA from whole ear skin tissue was extracted using RNeasy® Mini Kit (Qiagen), according to the manufacturer’s instructions. cDNA was synthesized using the PrimeScript RT Reagent Kit (Takara). Quantification of transcripts was performed on a Mx3005P QPCR system (Agilent Technologies) for the IL-1b expression assay, using Maxima SYBR green/ROX qPCR Master Mix (Thermo Fisher Scientific). Quantification of all other transcriptos was performed using the Luna® Universal qPCR Master Mix (New England BioLabs), on a CFX384™ Real-Time System (BIO-RAD). The following primers were used: *Il-1b* Forward: 5’TGTGCAAGTGTCTGAAGCAGC3’, Reverse: 5’TGGAAGCAGCCCTTCATCTT 3’. *Cd74* Forward: 5’ CACCACTGCTTACTTCCTGTACCAG 3’, Reverse: 5’GGTCATGTTGCCGTACTTGGTAACG 3’. *H2-aa* Forward: 5’ CAAGGTGGAGCACTGGGGC 3’, Reverse: 5’ CTGACATGGGGGCTGGAATCTCAG 3’. *RetnIa* Forward: 5’ GGCGTATAAAAGCATCTCATCTGGC 3’, Reverse: 5’GAGAGTCTTCGTTACAGTGGAGGG 3’. All samples were run in triplicates. qRT-PCR data are displayed as fold change compared to means of the respective control groups ± SD.

### Staining for flow cytometry

Preparation of single cell suspensions from skin and staining for extracellular markers were performed as described for Sorting of ear skin keratinocytes (see above). Antibodies used in different staining mixtures are anti-mouse CD45 (PerCP-Cy5.5 conjugated, clone 30-F11, eBioscience), anti-mouse CD326 (EpCAM) (APC conjugate, clone G8.8, eBioscience), anti-mouse γδTCR (FITC conjugated, clone eBioGoH3, eBioscience), anti-mouse CD86 (APC/Cy7 conjugated, clone GL1, Biolegend), anti-mouse CD3e (PE/Cy7 conjugated, clone 145-2C11, eBioscience), anti-mouse FcεRI (APC-conjugated, clone MAR-1, Invitrogen), anti-mouse Siglec-F (PE conjugated, clone E50-2440, BD Biosciences), anti-mouse Gr-1 (PerCP-Cy5.5 conjugated, clone RB6-8C5, eBioscience), anti mouse CD117 (biotin conjugated, clone 2B8, eBioscience), anti-mouse CD25 (PE conjugated, clone PC61.5, eBioscience), anti-mouse CD278/ICOS (PE/Cy7 conjugated, C398.4A, Biolegend), anti-mouse CD127/IL-7Ra (APC conjugated, clone SB-199, Biolegend), anti-mouse CD90.2 (Super Bright 780, clone 30-H12, Invitrogen), anti-mouse CD11c (biotin conjugated, clone N418, eBioscience). For the detection of biotin conjugated antibodies, either SA-V500 (BD Biosciences) or SA-PE/Cy7 (Biolegend) were used. FlowJo (TreeStar, Ashland, Ore) was used for data analysis.

## OVA model and skin scoring

### Analysis of DO11.10^+^/4get transgenic T cell response to epicutaneous OVA immunization

DO11.10^+^/4get (Murphy et al., 1990) splenocytes were incubated with CD4 (L3T4) MicroBeads (Miltenyi) and separated using LS Columns (Miltenyi) by positive selection. Isolated T cells were labelled with Cell Proliferation Dye eFluor™ 670 (eBiosceince) according to the manufacturer’s instructions. 2×10^6^ labelled transgenic T cells were resuspended in 100 µl of PBS and adoptively transferred into recipient mice by i.v. injection. 24h later, mice were epicutaneously immunized by painting 200 µl of 1mg/ml OVA/PBS solution onto the shaved back. This was repeated 2 more times at daily intervals. 48h after the last treatment, mice were sacrificed and single cell suspensions of inguinal LNs obtained by crushing the LNs between two glass slides, rinsing with 2ml of cold FACS buffer (0.5% BSA/2mM EDTA/PBS) and straining through 100 µM mesh. After centrifugation (1200 rpm, 5min, 41⍰C), cell pellets were resuspended in 75 µl of Fc blocking buffer (5 µg/ml Purified anti-mouse CD16/32 Antibody, Biolegend, in FACS buffer) and incubated on ice for 20 min. After pelleting, cells were resuspended in 75µl of FACS buffer containing anti-mouse CD4 (biotin conjugated, clone GK1.5, eBioscience) and anti-DO11.10 Ab (PE conjugated, clone KJ1-26, Miltenyi) and incubated on ice for 45 min.

Biotin detection was with SA-APC-Cy7 (Biolegend). After final washing, cells were resuspended in 200 µl of cold FACS buffer containing 1µg/ml DAPI. Samples were analysed on a BD FACSAria™ III instrument. FlowJo (TreeStar, Ashland, Ore) was used for data analysis.

### Quantification of total and antigen-specific IgE

Total mouse serum IgE levels were measured using the ELISA MAX™ Standard Set Mouse IgE kit (BioLegend) according to manufacturer’s instructions. All individual standards and mouse serum samples were analysed in triplicates, with means of triplicates displayed in graphs. For analysis of OVA-specific serum IgE, 96-well MaxiSorp ELISA plates (Nunc, Roskilde, Denmark) were coated with 2µg/ml of OVA solution in carbonate buffer and ELISA was performed using the ELISA MAX™ Standard Set Mouse IgE kit (BioLegend).

### Skin microbiome analysis

DNA was extracted from cheek skin swab samples using the QIAamp® DNA Mini Kit (QIAGEN GmbH, Hilden, Germany) as specified by the manufacturer for isolation of DNA from Gram-positive bacteria with modifications. 180 µL lysozyme (20 mg/mL) was used during for first lysis at 37°C for 30 min. 20 µL of proteinase K and 200 µL of buffer AL and lysis for 10 min omitting the 15 min incubation at 95°C. DNA quantity and quality were analyzed using a NanoDrop 1000 spectrophotometer (Thermo Fisher Scientific Germany BV & Co KG, Braunschweig, Germany). Negative controls were performed for each extraction batches. DNA was amplified using universal bacterial primers 515F and 806R targeting the V4 region of the 16S rRNA gene ((PMID: 22402401)). The primers were modified to include a unique barcode and Illumina primer sequences (P5 and P7). PCR was performed as previously described, including negative control and positive controls (mock community, HM-782D) (Schoilew et al., 2019). Libraries were prepared by PCR with 5 cycles using home-made primers combining the Illumina sequencing primers and the Illumina sequencing adapters and then purified using Agencourt AMPure XP beads (Beckman Coulter, Germany) following the manufacturer’s instructions. Purified products were checked for quality and concentration using Quant-iT™ PicoGreen® dsDNA Assay Kit (ThermoFisher scientific GmbH, Dreieich, Germany) and Qiaxcel instrument (QIAGEN GmbH, Hilden, Germany). An equimolar mix of all the PCR products was then sequenced on an Illumina® MiSeq instrument (V3 chemistry).

### Whole genome sequencing

High molecular weight genomic DNA (gDNA) was extracted from snap-frozen spleen of *Flg*^*ft/ft*^ BALB/c congenics (Saunders et al., 2016) and BALB/c *Flg*^*-/-*^ mice following the Animal Tissue DNA Isolation Kit (bionano genomics) following the manufacturer’s protocol for soft tissues. In brief, spleen tissue was homogenized and fixed on ice in a tissue grinder, followed by embedding in agarose plugs and proteinase K and RNase treatment of the plugs. Genomic DNA was recovered from plugs by agarase treatment and purified by drop dialysis against 1x TE buffer. Integrity of high molecular weight gDNA was determined by pulsed-field gel electrophoresis using the Pippin Pulse™ device (Sage Science). Most of the gDNA was larger than 300 kb in length. The gDNA was further purified with 1x volume AMPure XP beads (Beckman Coulter) following the manufacturer’s instructions. Long insert libraries were prepared using the SMRTbell Express Template Prep Kit 2.0 (Pacific Biosciences) following the manufacturer’s instructions. In brief, gDNA was sheared using the MegaRuptor™ device (Diagenode). 10 µg (*Flg*^*ft/ft*^ BALB/c congenic) and 18 µg (BALB/c *Flg*^*-/-*^) of sheared gDNA were used for PacBio SMRTbell™ library preparation. The libraries were subjected to size selection (about 15 kb) using the BluePippin™ instrument (Sage Science). Fractions were purified with 0,6x AMPure XP beads, and checked for their final size on the fragment analyzer (Agilent). The size selected library was loaded with 50 and 55 pM on plate on two Sequel SMRT cells (8M) each, sequencing primer v2 was used with the Sequel polymerase 2.0 and the Sequel sequencing kit 2.0, pre-extension time was 2 hours and run time was 30 hours for all SMRT cells. A total of 3 SMRT cells was sequenced for the *Flg*^*ft/ft*^ BALB/c congenics leading to a total yield of 43,6 Gb ccs reads and a total of 2 SMRT cells has been sequenced for the BALB/c *Flg*^*-/-*^ strain with a total yield of 41.4 Gb circular consensus sequences resulting in 18X genome coverage for both strains.

We used PacBio’s command line tool ccs (version 4.2.0, with default parameter, (https://github.com/PacificBiosciences/ccs)), that takes multiple subreads from the same SMRTbell molecule, to produce highly accurate long reads (HiFi reads). HiFi reads were generated for the three SMRT cells of the *Flg*^*ft/ft*^ BALB/c congenic strain and the two SMRT cells for the BALB/c *Flg*^*-/-*^ strain. The HiFi yield for the *Flg*^*ft/ft*^ BALB/c congenic strain was 45.7Gb (average read length 15.3Kb, coverage 16.2X), and 41.4 Gb (average read length 13.9Kb, coverage 14.7X) for the BALB/c *Flg*^*-/-*^ strain. Both HiFi read sets where assembled separately using the de novo assembler hifiasm with default parameters (version 0.7, https://github.com/chhylp123/hifiasm, https://arxiv.org/abs/2008.01237). The *Flg*^*ft/ft*^ BALB/c congenic strain assembly has a total size of 2.62Gb with N50 of 6.12Mb and 1373 contigs. The *Flg*^*-/-*^ BALB/c strain assembly has 1082 contigs with N50 of 6.5Mb and a total size of 2.59Gb. As the congenic region is located on chromosome 3, the variation analysis was limited to contigs corresponding to chromosome 3. We mapped all contigs to the mouse reference genome (GRCm38.p6) with minimap2 and separated the chromosome 3 contigs, resulting in 38 contigs. Gene annotation was performed by mapping the annotated mouse coding genes to the new assemblies. We generated liftOver chains by computing pairwise alignment chains to the mouse GRCm38 assembly using lastz (alignment parameters K=6000, L=8000, Y=3000, H=2000, default scoring matrix), axtChain (linearGap=loose, otherwise default parameters), chainNet and netChainSubset, as implemented in the UCSC genome browser script doBlastzChainNet.pl. We downloaded mouse GENCODE V25 genes from the UCSC genome browser table wgEncodeGencodeCompVM25 and used liftOver (https://genome.ucsc.edu/cgi-bin/hgLiftOver, parameter minMatch=0.8) to map the genes. For the variation analysis all HiFi reads (BALB/c *Flg*^*-/-*^ strain and *Flg*^*ft/ft*^ BALB/c congenic strain) were mapped to the BALB/c *Flg*^*-/-*^ assembly using pbmm2 (version 1.2.1, https://github.com/PacificBiosciences/pbmm2/). To reduce ambiguous mappings, the alignments were further filtered by restricting a maximum number of soft-clipped bases to 1. This resulted in two alignment files – one for the *Flg*^*ft/ft*^ BALB/c congenic reads and one for the BALB/c *Flg*^*-/-*^ reads. The HiFi mapped reads of the 38 chromosome 3 contigs were used for the variant analysis. We used freebayes (version v1.3.2-46-g2c1e395, https://github.com/ekg/freebayes, https://arxiv.org/abs/1207.3907) to detect small polymorphisms and pbsv (version: 2.3.0, https://github.com/pacificbiosciences/pbsv/) for structural variations. We filtered variants based on a quality score QUAL>20 to detect robust variants and those unique to reads of the *Flg*^*ft/ft*^ BALB/c congenic strain to remove any sequencing or alignment errors. Finally, the Ensembl Variant Effect Predictor (VEP, version: 100.3, (McLaren et al., 2016)) was used to annotate variants predicted to have phenotypic impact based on the gene annotations. Using the filtered variant files and custom annotations (see above), we then listed those with either Deleterious or Potentially damagingpredicted effects. These were then inspected manually.

## Supplementary methods

Pseudocode for programs that were used:

### Circular consensus sequence

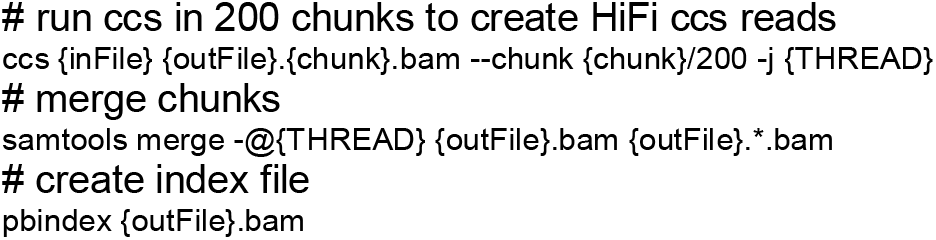

### De novo assembly

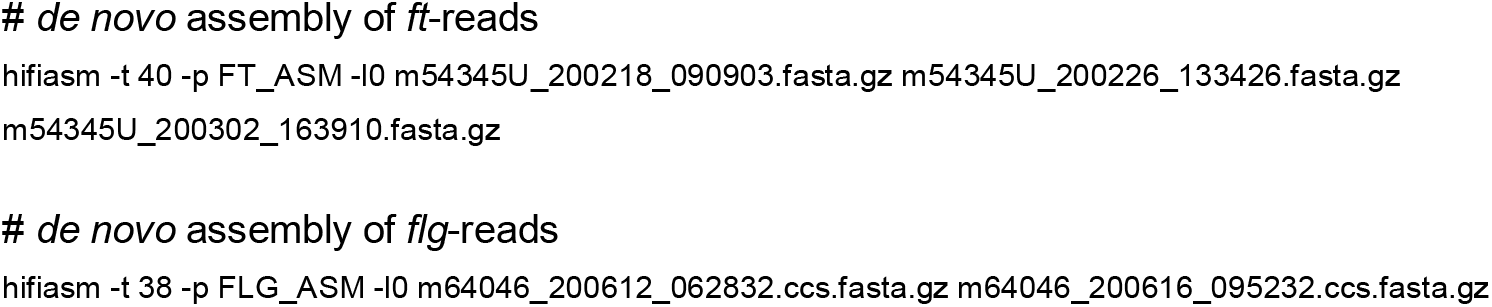

### Gene annotation

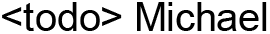

### Variation analysis

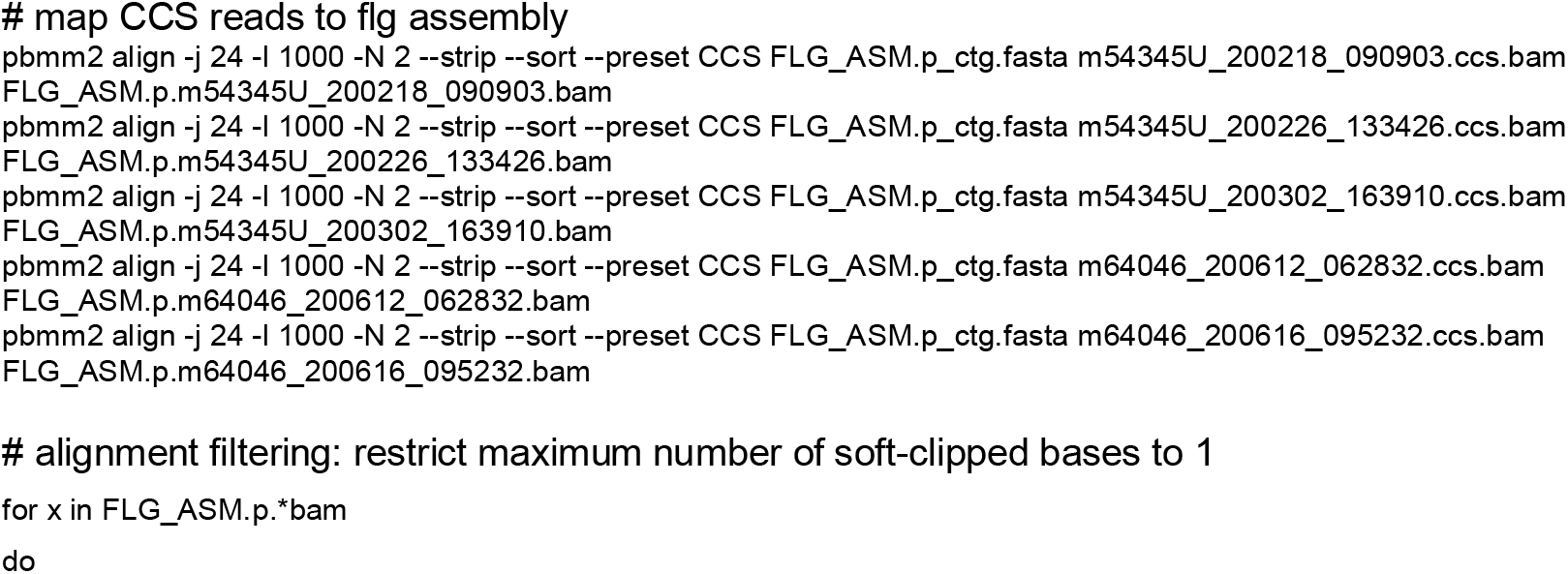
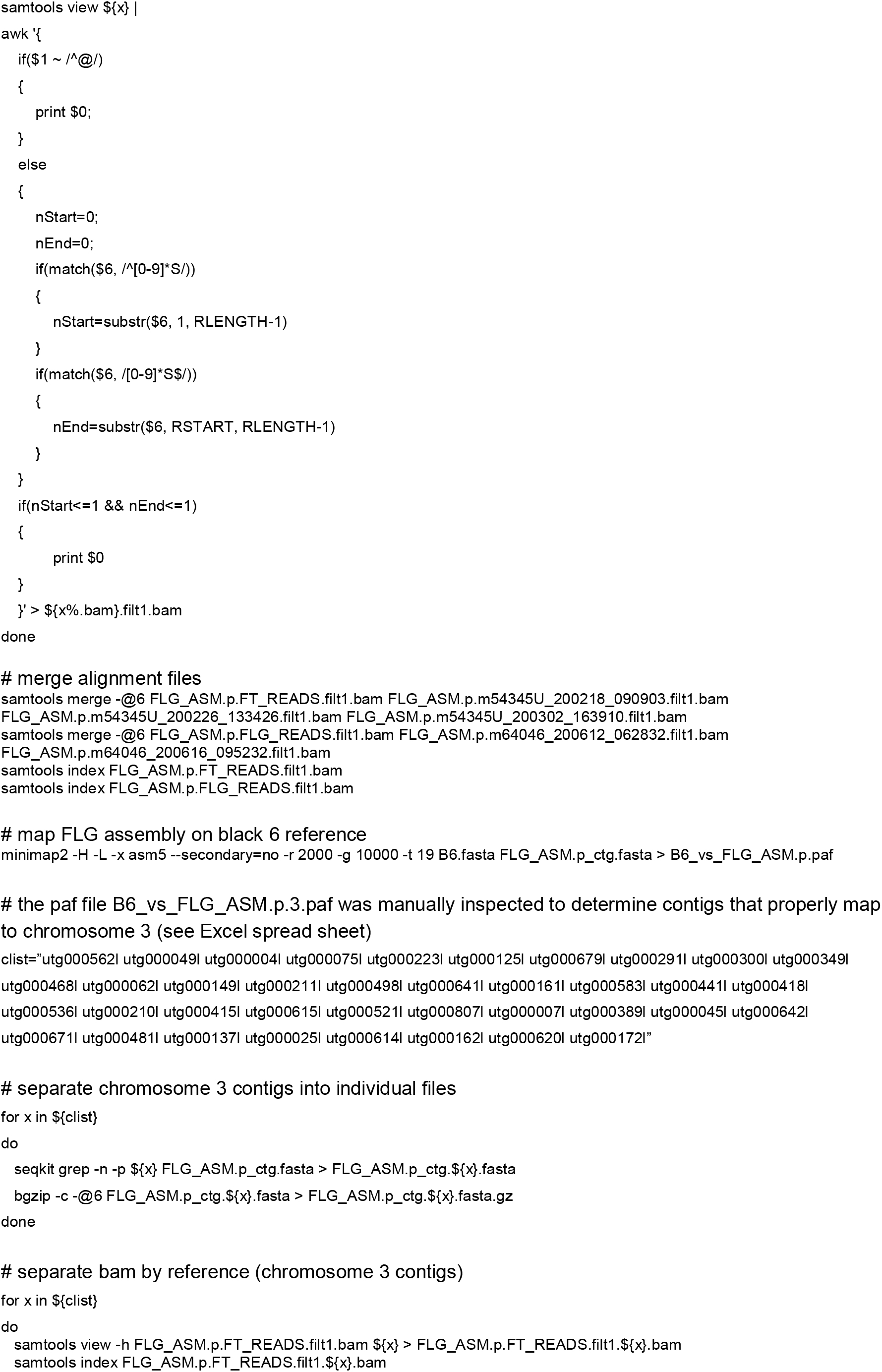
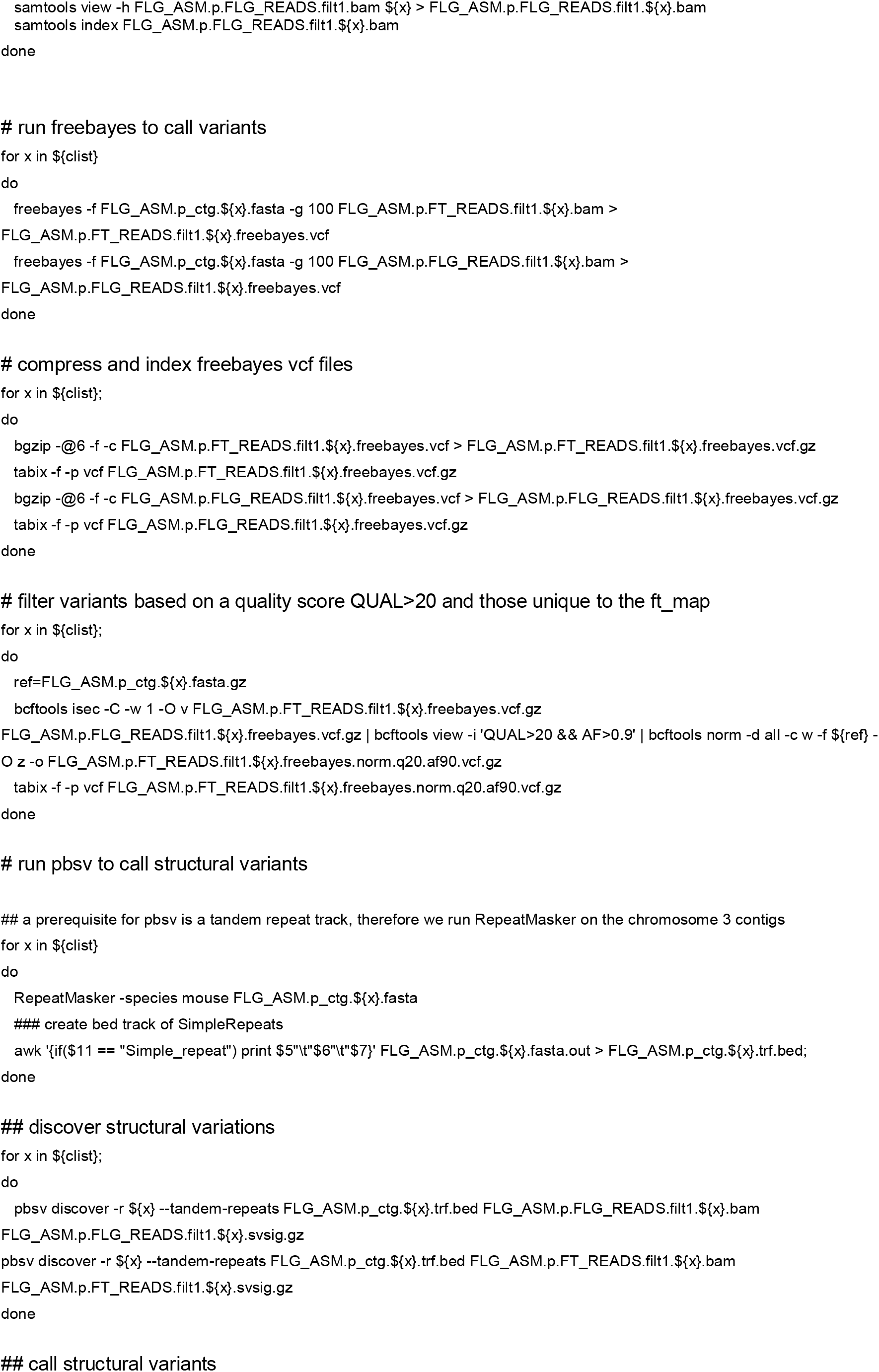
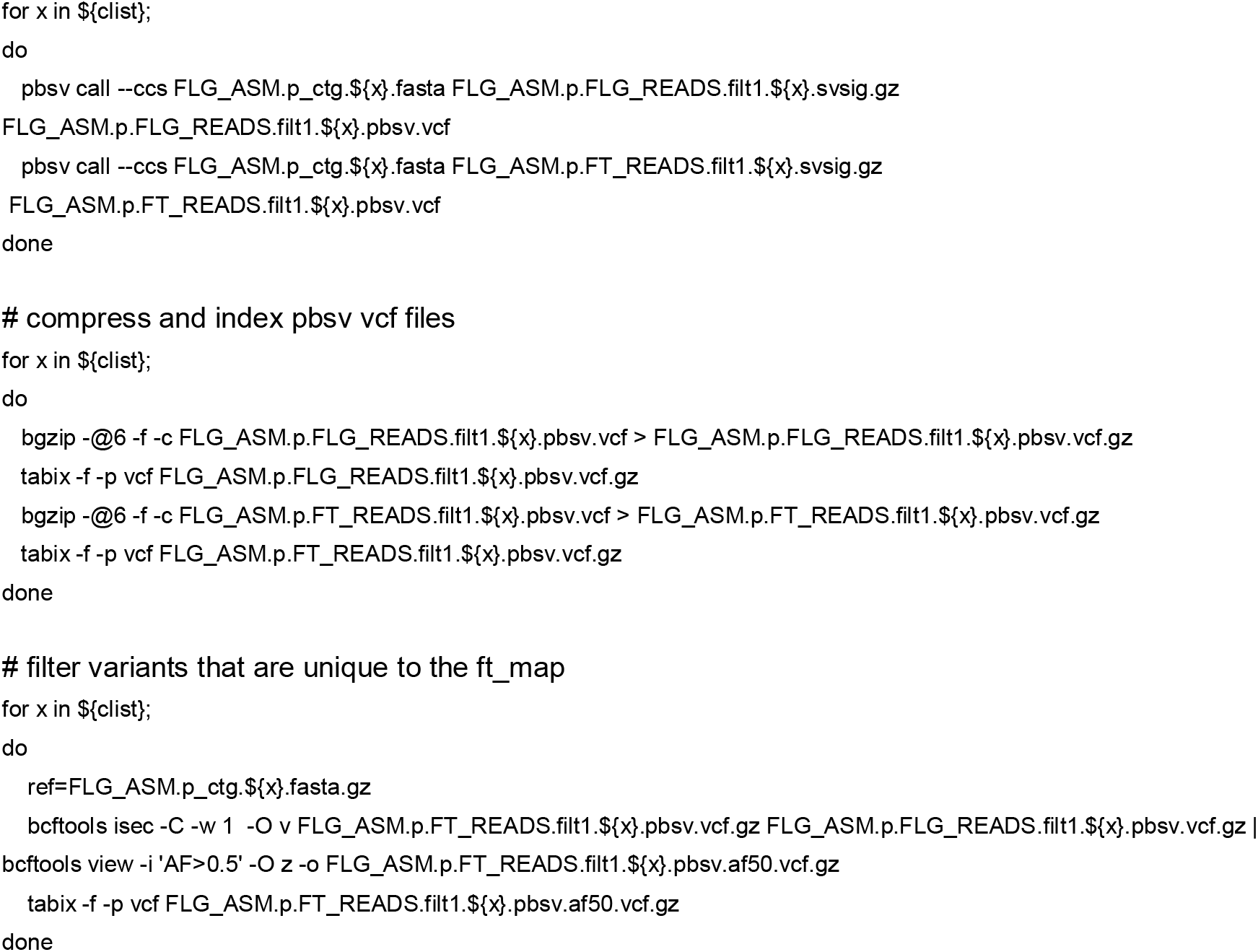

### Effect Prediction

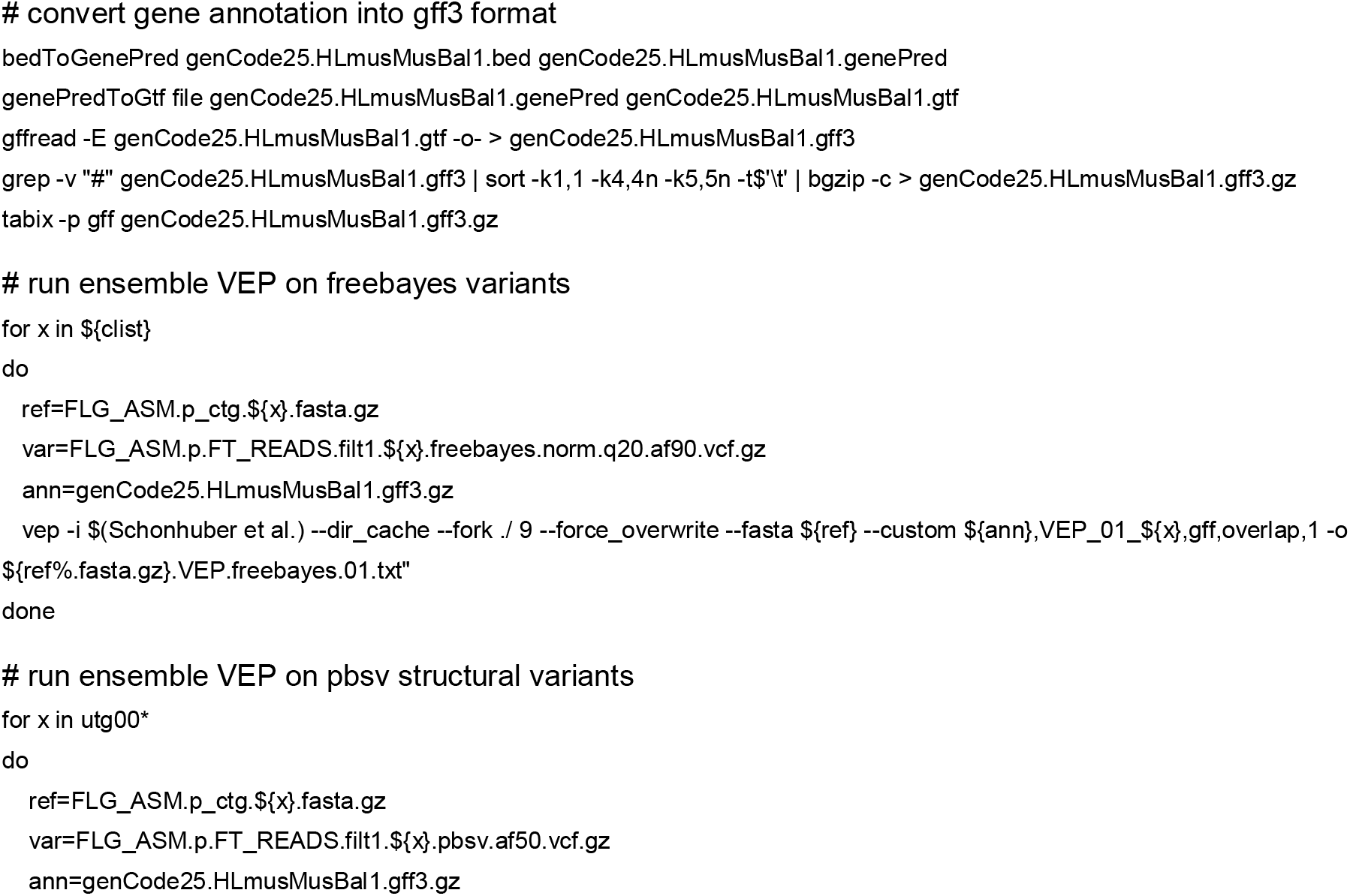
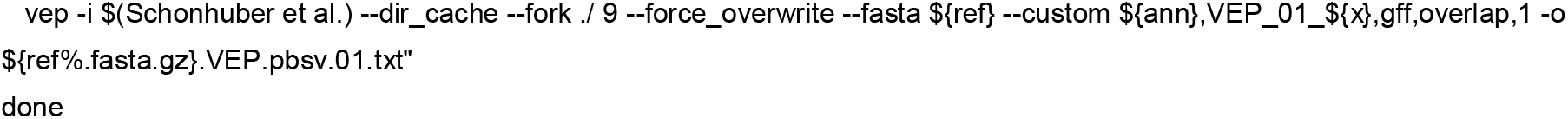

